# SV40 Large T antigen inhibits the host serine protease FAM111A through a zinc-dependent, cleavage-avoiding mechanism

**DOI:** 10.64898/2026.08.07.743322

**Authors:** Allison L. Welter, Sudhaker Dharavath, Yuka Machida, Julia R. Alvey, Sowmiya Palani, Morgan E. Crewe, Anh T. Q. Cong, Connor P. Jewell, Kosei Iwata, Lisa M. Jenkins, Matthew J. Schellenberg, Yuichi J. Machida

**Affiliations:** Developmental Therapeutics Branch, Center for Cancer Research, National Cancer Institute, Bethesda, MD, USA; Mayo Clinic Graduate School of Biomedical Sciences, Mayo Clinic, Rochester, MN, USA; Department of Molecular Pharmacology and Experimental Therapeutics, Mayo Clinic, Rochester, MN, USA; Department of Oncology, Division of Oncology Research, Mayo Clinic, Rochester, MN, USA; Department of Biochemistry and Molecular Biology, Mayo Clinic, Rochester, MN, USA; Laboratory of Cell Biology, Center for Cancer Research, National Cancer Institute, Bethesda, MD, USA

## Abstract

SV40 Large T antigen (LT) is essential for viral replication and a key determinant of host range. This host-range function is mediated by the C-terminal domain (LT-C) through binding to the host serine protease FAM111A, but the underlying mechanism has remained unclear. Here, we report the X-ray crystal structure of the FAM111A serine protease domain in complex with LT-C, revealing the structural basis for direct inhibition of FAM111A. LT-C uses a previously unrecognized zinc-binding motif and a P1-like phenylalanine residue to engage the FAM111A active site through a substrate-mimicking mechanism while avoiding proteolytic cleavage and covalent complex formation. Mutations disrupting either feature abolish FAM111A inhibition and impair SV40 propagation in cells. Consistent with this mechanism, SV40 host restriction requires FAM111A protease activity, which must be antagonized by LT-C for productive infection. Together, these findings define a zinc-dependent, cleavage-avoiding mechanism of protease inhibition that highlights an evolutionary arms race between SV40 and host antiviral proteases.

## Introduction

Host-virus interactions reflect an ongoing evolutionary arms race in which host cells deploy antiviral defenses to restrict viral infection, while viruses evolve countermeasures to overcome these barriers and enable productive replication. A powerful approach to dissecting these interactions is the study of host-range mutant viruses, which lack viral factors required to counteract specific host defense mechanisms. Analyses of host-range mutants have revealed key antiviral pathways in diverse virus families, including orthopoxviruses and polyomaviruses^1,2^, making them a valuable entry point for uncovering host restriction mechanisms and the viral strategies that overcome them.

Polyomaviruses are nonenveloped double-stranded DNA viruses with small genomes of approximately 5 kb that replicate in the nucleus of host cells. Simian virus 40 (SV40) is a macaque polyomavirus and the most extensively studied member of this family, serving as a model system for studies of DNA replication, transcription, and oncogenic transformation^3,4^. SV40 expresses a number of viral proteins, including early proteins small T antigen (sT) and large T antigen (LT), and late proteins VP1, VP2, and VP3, and agnoprotein. LT is a multifunctional protein, playing a central role in viral DNA replication by functioning as a helicase^5,6^ and in transcriptional regulation of viral gene expression^7,8,9,10^, while reprograming the host cell cycle through interactions with host cell proteins such as p53 and retinoblastoma protein (Rb)^11,12^.

In addition to its roles in DNA replication and transcription, SV40 LT contributes to host range through its C-terminal region (LT-C; amino acids 627-708), which is essential for productive infection^13,14,15,16^. Genetic studies identified LT-C as the host-range domain by showing that deletions or frameshift mutations within this region generate host-range mutants that replicate efficiently in permissive cells but fail to establish productive infection in non-permissive cells. Interestingly, despite relatively high overall conservation of Large T among polyomaviruses, LT-C is the most sequence-divergent region of the protein. Moreover, the LT-C host-range domain is present in only a subset of polyomaviruses, including SV40, JC virus, BK virus, and SA12^17^, suggesting that it represents a specialized evolutionary adaptation rather than a core viral function.

A proteomics study identified the protease domain of FAM111 trypsin-like peptidase A (FAM111A) as a prominent interactor of LT-C^1^. In the same study, FAM111A was shown to restrict SV40 host-range mutant infection in non-permissive cells, as depletion of FAM111A permits viral propagation. FAM111A was also found to localize at SV40 replication centers^18^, supporting its role as a host restriction factor. These observations placed FAM111A at the center of a long-standing host-range phenotype but did not explain how the small LT-C domain counteracts FAM111A-mediated antiviral restriction. FAM111A also functions as an antiviral factor against orthopoxviruses. Poxviruses lacking the serine protease inhibitor SPI-1 exhibit severe replication defects in non-permissive cells, which can be rescued by depletion of FAM111A^2^. Our previous study showed that SPI-1 directly inhibits FAM111A through a classical serpin mechanism involving covalent complex formation, demonstrating that direct inhibition of FAM111A protease activity is an effective strategy for overcoming host restriction^19^.

FAM111A is a serine protease that contains an N-terminal PCNA-interacting peptide (PIP) box^20^, two ubiquitin-like (UBL) domains^21^, and a C-terminal serine protease domain (SPD)^22,23^. Through its interaction with PCNA, FAM111A localizes to DNA replication forks^20^, where it promotes DNA replication at protein obstacles, including persistent topoisomerase cleavage complexes and trapped poly(ADP-ribose) polymerases (PARP)-DNA complexes^24^. We recently showed that FAM111A exhibits chymotrypsin-like peptidase activity and preferentially cleaves substrates containing phenylalanine at the P1 position, the residue immediately N-terminal to the scissile bond^22^. Structural analysis further revealed a narrow active-site cleft with a hydrophobic S1 specificity pocket that accommodates P1 phenylalanine residues, providing the structural basis for this substrate specificity^22^.

Although LT-C has long been genetically linked to SV40 host range through antagonism of FAM111A, how this small viral region neutralizes FAM111A has remained unknown. Here, we show that the C-terminal region of SV40 Large T antigen functions as a direct inhibitor of FAM111A and determine the crystal structure of the LT-C:FAM111A protease complex. The structure reveals that LT-C inhibits FAM111A protease activity through a zinc-dependent, cleavage-avoiding mechanism without undergoing proteolytic cleavage or covalent complex formation. We further identify a P1-like phenylalanine residue and a previously unrecognized zinc-binding motif within LT-C that are essential for FAM111A inhibition. Together, these findings establish LT-C as a viral serine protease inhibitor domain, define a distinct zinc-dependent mechanism for uncoupling substrate recognition from proteolysis, and reveal protease inhibition as a viral counter-defense mechanism that expands host-range.

## Results

### SV40 Large T antigen’s C-terminus directly inhibits FAM111A protease activity

To test whether LT-C directly inhibits FAM111A protease activity, we examined the effect of LT-C on the purified FAM111A SPD using our previously established *in vitro* peptidase assay^22^. Recombinant FAM111A SPD (wild-type and active site mutant S541A), GST, and GST-LT-C were expressed and purified, with LT-C consisting of the host-range region of Large T (amino acids 627-708; Fig. 1A and Supplementary Fig. 1A). GST-LT-C inhibited FAM111A SPD activity in a dose-dependent manner, whereas GST alone had no effect (Fig. 1B and Supplementary Fig. 1B), demonstrating that LT-C acts as a FAM111A inhibitor. Consistent with direct inhibition, FAM111A SPD and GST-LT-C interacted in GST pull-down assays, and this interaction was also observed with the S541A mutant (Fig. 1C), indicating that binding is independent of the active site serine. These results suggest that LT-C is a direct inhibitor of FAM111A protease activity.

**Figure 1.**
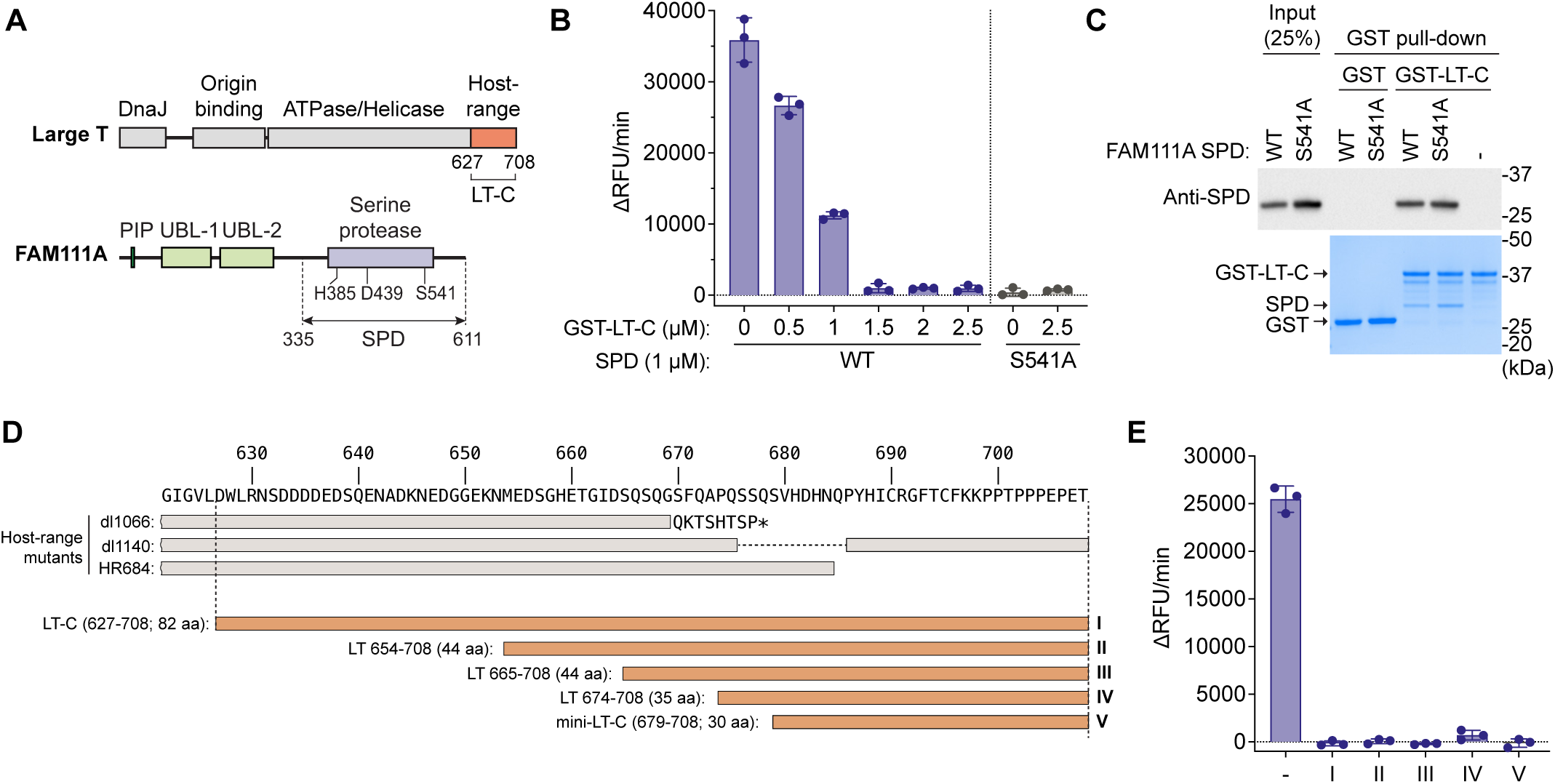
The C-terminal region of SV40 Large T antigen directly inhibits FAM111A protease activity. **(A)** Schematic domain organization of SV40 Large T and FAM111A. The C-terminal host-range region of Large T (residues 627-708; LT-C) used in this study is indicated in orange. The serine protease domain (SPD) of FAM111A and the catalytic triad (H385, D439, and S541) are indicated. PIP: PCNA-interacting peptide; UBL: ubiquitin-like. **(B)** Peptidase activity of FAM111A SPD measured using the fluorogenic substrate Suc-AAPF-AMC in the presence of increasing concentrations of GST–LT-C, expressed as the change in relative fluorescence units per minute (ΔRFU/min). Reactions without GST-LT-C contained GST as a control. Values are mean ± SD of three technical replicates. **(C)** In vitro GST pull-down assay of GST-LT-C incubated with FAM111A SPD, either wild-type (WT) or active site mutant (S541A). Pull-down of FAM111A SPD was detected by western blotting with anti-FAM111A SPD antibody (top), and GST proteins were visualized by SDS-PAGE followed by Coomassie staining (bottom). The positions of GST, GST-LT-C, and FAM111A SPD are indicated on the left, and molecular weight markers are shown on the right. **(D)** Schematic representation of LT-C truncation constructs used to map the region required for FAM111A inhibition. Colored bars (I-V) indicate recombinant GST-LT-C fragments tested in this study. Grey bars represent previously characterized SV40 Large T host-range mutants (dl1066, dl1140, and HR684) and are shown to indicate regions previously implicated in host-range function. Amino acid positions are indicated above the sequence. **(E)** *In vitro* peptidase activity of FAM111A SPD measured as in (B) in the presence of GST-LT-C truncation constructs as indicated in (D). Values are mean ± SD of three technical replicates.

To determine the region within LT-C responsible for FAM111A inhibition, we generated a series of truncation mutants spanning the C-terminal region of Large T (Fig. 1D, Supplementary Fig. 1C). Progressive deletion of LT-C N-terminal sequences, informed by regions previously implicated in host-range mutants^14,16^, revealed that constructs encompassing residues 664-708, 665-708, 674-708, and 679-708 all retained inhibitory activity comparable to full-length LT-C (Fig. 1E, Supplementary Fig. 1D). Notably, a minimal fragment corresponding to residues 679-708 (hereafter mini-LT-C) was sufficient to inhibit FAM111A SPD activity to the same extent as longer constructs, mapping the inhibitory activity of LT-C to the C-terminal 30 amino acids of Large T antigen.

### Structural basis for LT-C-mediated inhibition of FAM111A

Because Large T antigen has not previously been recognized as a protease inhibitor, we sought to determine the molecular basis of LT-C-mediated FAM111A inhibition using X-ray crystallography. Recombinant FAM111A SPD was incubated with a mini-LT-C peptide produced in *E. coli*, and the resulting complex was purified by size exclusion chromatography. However, crystals obtained from this preparation contained only SPD with no detectable mini-LT-C, likely because a strong crystal lattice packing occluded the protease active site. We therefore used a dimerization-defective SPD variant containing V347D and V351D substitutions (hereafter SPD^VDVD^; Supplementary Fig. 2A)^22^ which fortuitously crystallized in a P2_1_2_1_2 lattice that did not occlude the active-site. The apo SPD^VDVD^ structure was determined at 1.86 Å resolution with a single molecule in the asymmetric unit (Protein Data Bank (PDB) IDs: XXXX; Supplementary Table 1 and Supplementary Fig. 2B). Although SPD^VDVD^ lacks protease activity because it does not dimerize effectively in solution^22^, the crystal lattice generated a symmetry-related dimer through a two-fold crystallographic axis. This dimer closely resembled wild-type SPD, with only slight distortion relative to the wild-type dimer and with the active site and oxyanion hole maintained in an active conformation. Consistent with this overall similarity, the SPD^VDVD^ structure superimposed closely on the previously reported wild-type SPD structure^22^ (Cα root-mean-square deviation (RMSD) = 0.35 Å over 192 Cα atoms), with differences largely restricted to flexible loop regions (Supplementary Fig. 2C).

The SPD^VDVD^:mini-LT-C complex was purified by size-exclusion chromatography as a single peak consistent with a stable complex, and the corresponding fractions were confirmed by SDS-PAGE (Supplementary Fig. 3A). We next crystalized and determined the X-ray crystal structure of SPD^VDVD^:mini-LT-C complex at 1.66 Å resolution (PDB ID: XXXX; Supplementary Table 1). The complex crystallized in the P2_1_ space group, with two SPD^VDVD^:mini-LT-C complexes in the asymmetric unit exhibiting essentially identical conformations (Cα RMSD = 0.07 Å over 207 Cα atoms; Supplementary Fig. 3B,C). The structure revealed that mini-LT-C spans the substrate-binding cleft of SPD, forming an antiparallel β-sheet with the β10 strand while extending into the active site. Unexpectedly, the structure also revealed a zinc ion coordinated by mini-LT-C, a previously unrecognized feature of the inhibitory complex (Fig. 2A).

**Figure 2.**
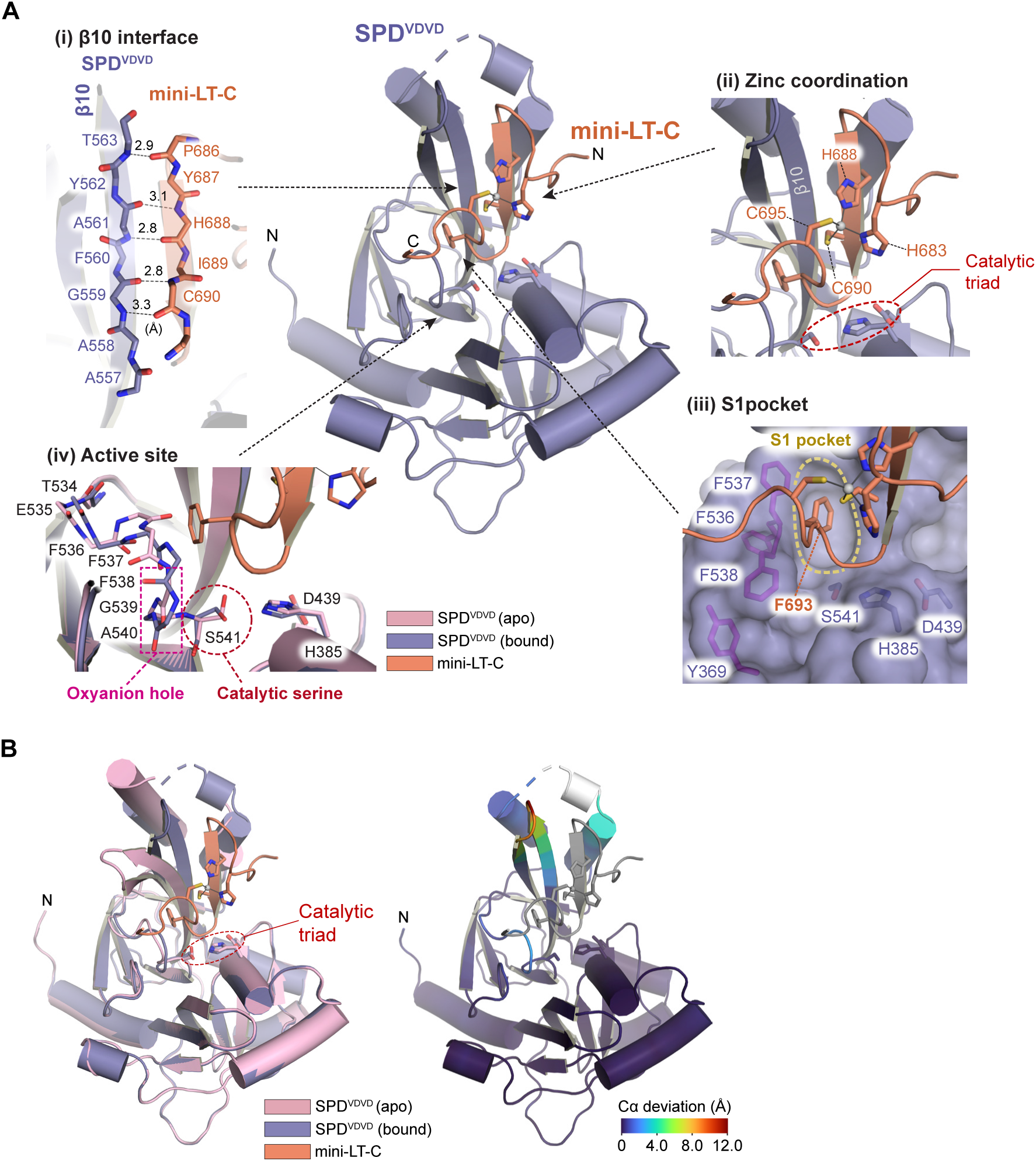
X-ray crystal structure of the FAM111A SPD^VDVD^:mini-LT-C complex. **(A)** Overall structure of the SPD^VDVD^:mini-LT-C complex. SPD^VDVD^ is shown as a purple cartoon and mini-LT-C as an orange cartoon. Insets highlight the major interaction interfaces between SPD and mini-LT-C: (i) the antiparallel β-sheet formed between the β10 strand of SPD and the N-terminal region of mini-LT-C, with backbone hydrogen bonds indicated as dashed lines and distances (Å); (ii) the zinc-binding site coordinated by His683, His688, Cys690, and Cys695 of mini-LT-C; (iii) insertion of the P1-like residue Phe693 into the S1 pocket of SPD; and (iv) comparison of the active site in the apo (pink) and mini-LT-C-bound (purple) SPD^VDVD^ structures, highlighting conformational changes in the oxyanion-hole loop and catalytic Ser541. **(B)** Superposition of the apo SPD^VDVD^ (pink) and mini-LT-C-bound SPD^VDVD^ (purple) structures. Left, structural overlay showing the overall similarity between the two structures. Right, Cα deviation mapped onto the bound structure, with colors indicating the magnitude of local conformational changes.

The N-terminal region of mini-LT-C is anchored to SPD through an antiparallel β-sheet formed with the β10 strand of the protease domain (Fig. 2A,i). The β-sheet is stabilized primarily by backbone hydrogen bonds between LT-C residues Pro686-Cys690 and SPD residues Gly559-Thr563, positioning the C-terminal region of LT-C to extend into the catalytic site. Inspection of the LT-C structure revealed that His683, His688, Cys690, and Cys695 form a tetrahedral zinc-binding site. Strong electron density at this site was assigned as a zinc ion based on its coordination geometry and anomalous diffraction signal (Fig. 2A,ii and Supplementary Fig. 3D). To our knowledge, this zinc-binding motif has not previously been described in polyomavirus Large T antigens. The C-terminal region of mini-LT-C extends into the substrate-binding cleft, where the P1-like residue Phe693 inserts its aromatic side chain deeply into the S1 specificity pocket of SPD (Fig. 2A,iii). Consistent with our previous finding that FAM111A preferentially recognizes phenylalanine at the P1 position, the S1 pocket is lined by Tyr369, Phe536, Phe537, and Phe538, forming a predominantly hydrophobic environment that accommodates the P1-like phenylalanine side chain. Superposition of the apo and LT-C-bound structures demonstrated that the overall fold of the protease domain is highly conserved, with an RMSD of 0.6 Å over 260 Cα atoms (Fig. 2B). Nevertheless, LT-C binding induced localized conformational changes in the active-site loop comprising residues Glu535-Ser541, which contributes to both the S1 pocket and the oxyanion hole (Fig. 2A,iv and Fig. 2B). Accompanying these changes, the side chain of the catalytic Ser541 rotates relative to the apo structure, with well-defined electron density supporting the assigned conformations (Supplementary Fig. 3E), orienting its hydroxyl group away from the catalytically competent configuration. These local rearrangements enable the P1-like residue Phe693 to occupy and occlude the S1 specificity pocket while repositioning the catalytic serine into a catalytically incompetent conformation.

### Zinc-binding motif in LT-C is required for interaction with and inhibition of FAM111A

To investigate the functional importance of the newly identified zinc-binding motif, we first asked whether LT-C binds zinc in solution. The structure predicts that His683, His688, Cys690, and Cys695 coordinate a zinc ion (Fig. 2A,ii). Consistent with this prediction, inductively coupled plasma mass spectrometry (ICP-MS) revealed that GST-LT-C purified from *E. coli* contained substantially higher levels of zinc than GST alone (Fig. 3A), confirming that recombinant LT-C retains bound zinc following purification.

**Figure 3.**
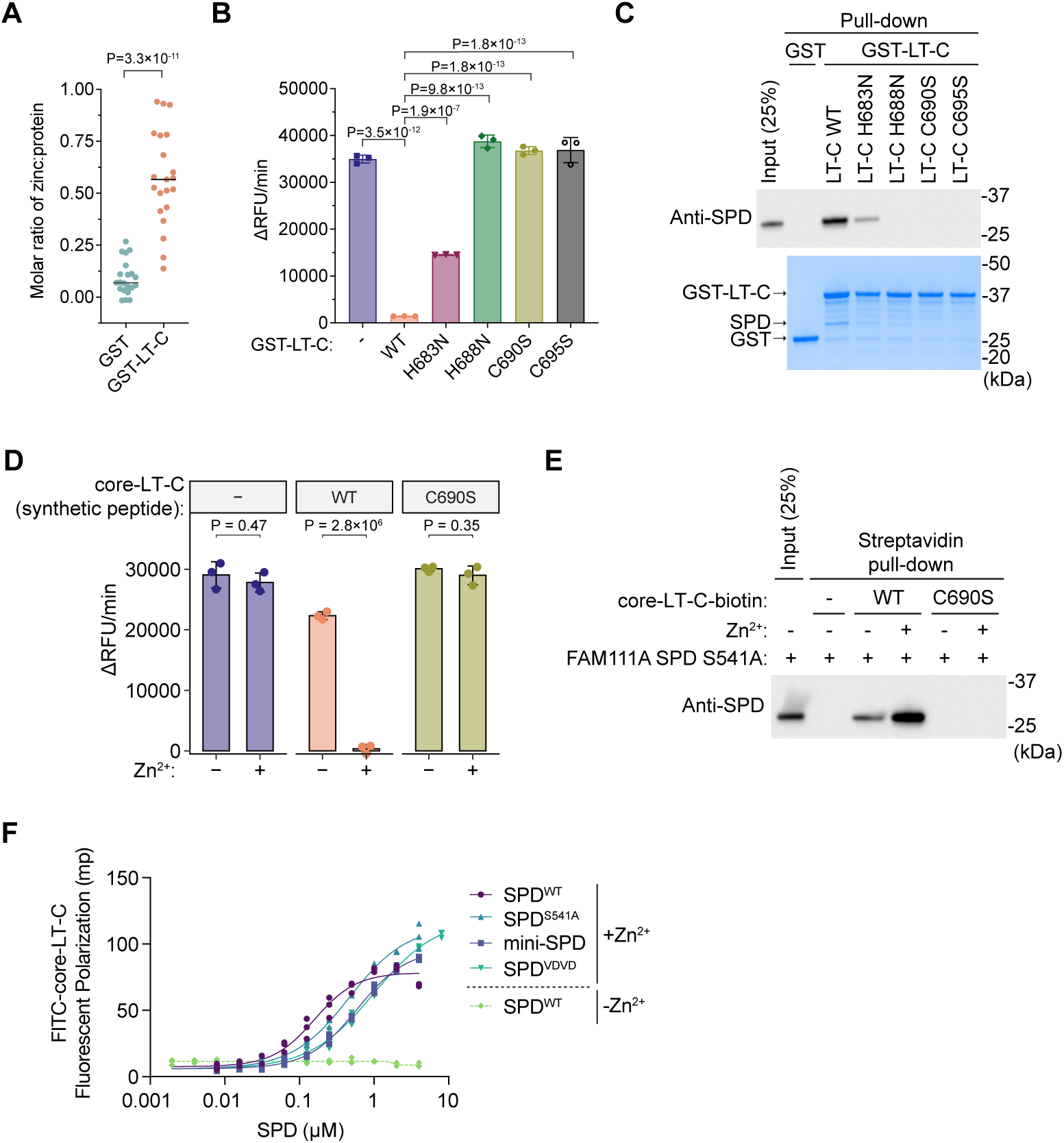
A zinc-binding motif is required for interaction with and inhibition of FAM111A. **(A)** Zinc content of purified GST and GST-LT-C proteins measured by Inductively coupled plasma mass spectrometry (ICP-MS), shown as the molar ratio of zinc to protein. Individual measurements are shown as points, medians indicated by black lines. Statistical significance was assessed using an unpaired two-tailed t-test. **(B)** *In vitro* peptidase activity of FAM111A SPD measured using Suc-AAPF-AMC as a substrate in the presence of GST-LT-C (wild-type or zinc-binding motif mutants H683N, H688N, C690S, and C695S). Reactions without GST-LT-C contained GST as a control. Values are mean ± SD of three technical replicates. Statistical significance was determined by one-way ANOVA with Dunnett’s multiple-comparison test. RFU: relative fluorescence units. **(C)** *In vitro* GST pull-down assay of FAM111A SPD with GST-LT-C (wild-type or zinc-binding motif mutants H683N, H688N, C690S, and C695S). Bound FAM111A SPD was detected by western blotting (top), and GST proteins were visualized by Coomassie staining (bottom). Molecular weight markers are shown on the right. **(D)** *In vitro* peptidase activity of FAM111A SPD measured in the presence of synthetic core-LT-C peptide (5 µM) corresponding to residues 679-704 (wild-type or C690S) incubated with or without ZnCl_2_ (2.5 µM). Assays were performed using Suc-AAPF-AMC as a substrate. Values are mean ± SD of three technical replicates. Statistical significance was determined by unpaired two-tailed t-tests. **(E)** *In vitro* streptavidin pull-down assay using biotinylated core-LT-C peptides corresponding to residues 679-704 (wild-type or C690S) incubated with FAM111A SPD S541A in the presence or absence of a twofold molar excess of ZnCl_2_. Bound FAM111A SPD was detected by western blotting. Molecular weight markers are shown on the right. **(F)** Binding of FITC-core-LT-C to the indicated SPD proteins was measured using fluorescence polarization anisotropy. Wild-type SPD was assayed either in the presence (+Zn^2+^) or absence (-Zn^2+^) of zinc, whereas the remaining SPD variants were assayed in the presence of Zn^2+^. Data were fit to a 4-parameter dose-response curve (solid and broken lines). Data shown from three independent experiments.

To determine whether the zinc-coordinating residues are required for LT-C-mediated inhibition of FAM111A, we generated recombinant GST-LT-C proteins in which His683 and His688 were individually substituted with asparagine and Cys690 and Cys695 with serine, substitutions predicted to disrupt zinc coordination while minimizing structural perturbation (Supplementary Fig. 4A). In both GST pull-down assays and FAM111A peptidase assays, three mutants (H688N, C690S, and C695S) exhibited little or no interaction with SPD and failed to inhibit its protease activity (Fig. 3B,C and Supplementary Fig. 4B). By contrast, H683N retained weak SPD binding and partial inhibitory activity, suggesting that zinc coordination may be partially maintained or compensated in this mutant. These findings identify His688, Cys690, and Cys695 as critical determinants of LT-C binding and inhibition, supporting a critical role for the zinc-binding motif in LT-C-mediated inhibition of FAM111A.

Because H683N retained weak SPD binding and partial inhibitory activity, we investigated whether nearby residues could compensate for the loss of His683. Inspection of the LT-C structure identified His681 adjacent to His683, and because aspartate residues can occasionally participate in zinc coordination^25^, we additionally examined Asp682 (Supplementary Fig. 4C). Whereas the H681N and D682A single mutants retained both SPD binding and inhibitory activity, the H681N/H683N double mutant exhibited markedly reduced binding and inhibition compared with H683N alone (Supplementary Fig. 4D-G). These findings indicate that His681 can partially compensate for the loss of His683, supporting a model in which His683, His688, Cys690, and Cys695 constitute the principal residues of a functional H2C2-like zinc-binding motif required for LT-C-mediated inhibition of FAM111A.

### Zinc is required for LT-C-mediated interaction with and inhibition of FAM111A

To directly test whether LT-C-mediated inhibition of FAM111A depends on zinc, we synthesized a 26-amino acid LT-C peptide encompassing residues 679-704 (hereafter referred to as core-LT-C), which contains the complete zinc-binding motif. In FAM111A SPD peptidase assays, the wild-type core-LT-C peptide exhibited little inhibitory activity in the absence of added zinc (Fig. 3D and Supplementary Fig. 5A). Addition of zinc, however, resulted in robust inhibition of FAM111A SPD, while zinc alone in the absence of LT-C peptide did not. By contrast, a core-LT-C peptide carrying the C690S substitution failed to inhibit FAM111A even in the presence of zinc, demonstrating that zinc-dependent inhibition requires an intact zinc-binding motif.

We next asked whether zinc similarly promotes LT-C binding to FAM111A. In peptide pull-down assays using C-terminally biotinylated core-LT-C peptides, zinc markedly enhanced the interaction between the wild-type peptide and SPD, whereas the C690S peptide failed to bind under either condition (Fig. 3E). Although weak binding of the wild-type peptide was detected in the absence of added zinc, this interaction was abolished by EDTA (Supplementary Fig. 5B), suggesting that trace zinc contamination in the reaction supported residual binding. Taken together, these findings demonstrate that zinc is required for both LT-C binding to and inhibition of FAM111A.

### LT-C binds FAM111A with high affinity

To quantify the affinity of the FAM111A SPD:LT-C interaction, we used a fluorescence polarization assay with a fluorescein-labeled core-LT-C peptide pre-complexed with zinc (Fig. 3F). Wild-type SPD bound the core-LT-C probe with high affinity, yielding a K_D_ of 158 ± 16 nM, consistent with the robust interaction observed in pull-down assays (Fig. 3F and Supplementary Fig. 5C). In contrast, no measurable binding was detected in the absence of zinc, demonstrating that LT-C binding to FAM111A is zinc-dependent, which is consistent with its structural role in the LT-C zinc-binding motif. The catalytically inactive S541A mutant also bound core-LT-C, although with a modestly reduced affinity (K_D_ = 470 ± 30 nM), supporting a mechanism in which LT-C binding does not require covalent complex formation with the active-site serine. We also examined two predominantly monomeric, catalytically inactive SPD variants^22^, mini-SPD and the SPD^VDVD^ which was co-crystalized with mini-LT-C. Both retained measurable binding to core-LT-C, with K_D_ values of 1320 ± 30 nM and 1010 ± 90 nM, respectively. These results are consistent with the largely preserved interaction interface observed in the monomeric and dimeric crystal structures, but indicate that LT-C preferentially binds the active, dimeric conformation of FAM111A. Taken together, these findings establish that LT-C engages FAM111A through a high-affinity, zinc-dependent interaction.

### The β10 interface and Phe693 are required for LT-C binding to and inhibition of FAM111A

The crystal structure identified two major interaction interfaces between LT-C and FAM111A SPD: an antiparallel β-sheet involving the β10 strand of the protease domain (Fig. 2A,i) and LT-C, and insertion of the P1-like residue Phe693 into the S1 specificity pocket (Fig. 2A,iii). To determine whether the β10 interface contributes to LT-C-mediated inhibition of FAM111A, we sought to disrupt the intermolecular β-sheet. Although the backbone atoms of His688 and Cys690 contribute to β-sheet formation (Fig. 2A,i), the side chains of these residues also coordinate the structural Zn^2+^ ion (Fig. 2A,ii). Consequently, mutating His688 or Cys690 would be expected to destabilize LT-C rather than selectively disrupt the β10 interface. We therefore targeted the complementary β10 strand in the FAM111A SPD by substituting Gly559 and Ala561 with proline (Fig. 4A, Supplementary Fig. 6A). Proline substitutions disrupt β-strand conformation through steric clash between the proline Cα and the adjacent carbonyl and have been successfully used to disrupt antiparallel β-strand interfaces, including the FAM111A-SPI-1 interaction^19,22^. In the absence of LT-C, the A561P substitution increased basal protease activity, whereas G559P exhibited a reduction relative to wild type (Fig. 4B, Supplementary Fig. 6B). Wild-type SPD was inhibited by mini-LT-C as expected, while the A561P or G559P mutants were not, indicating that disruption of the β10 interface impairs LT-C-mediated SPD inhibition. Consistent with these findings, both β10 mutants failed to bind GST-LT-C in pull-down assays (Fig. 4C), demonstrating that the β10 interface is required for stable LT-C association and protease inhibition.

**Figure 4.**
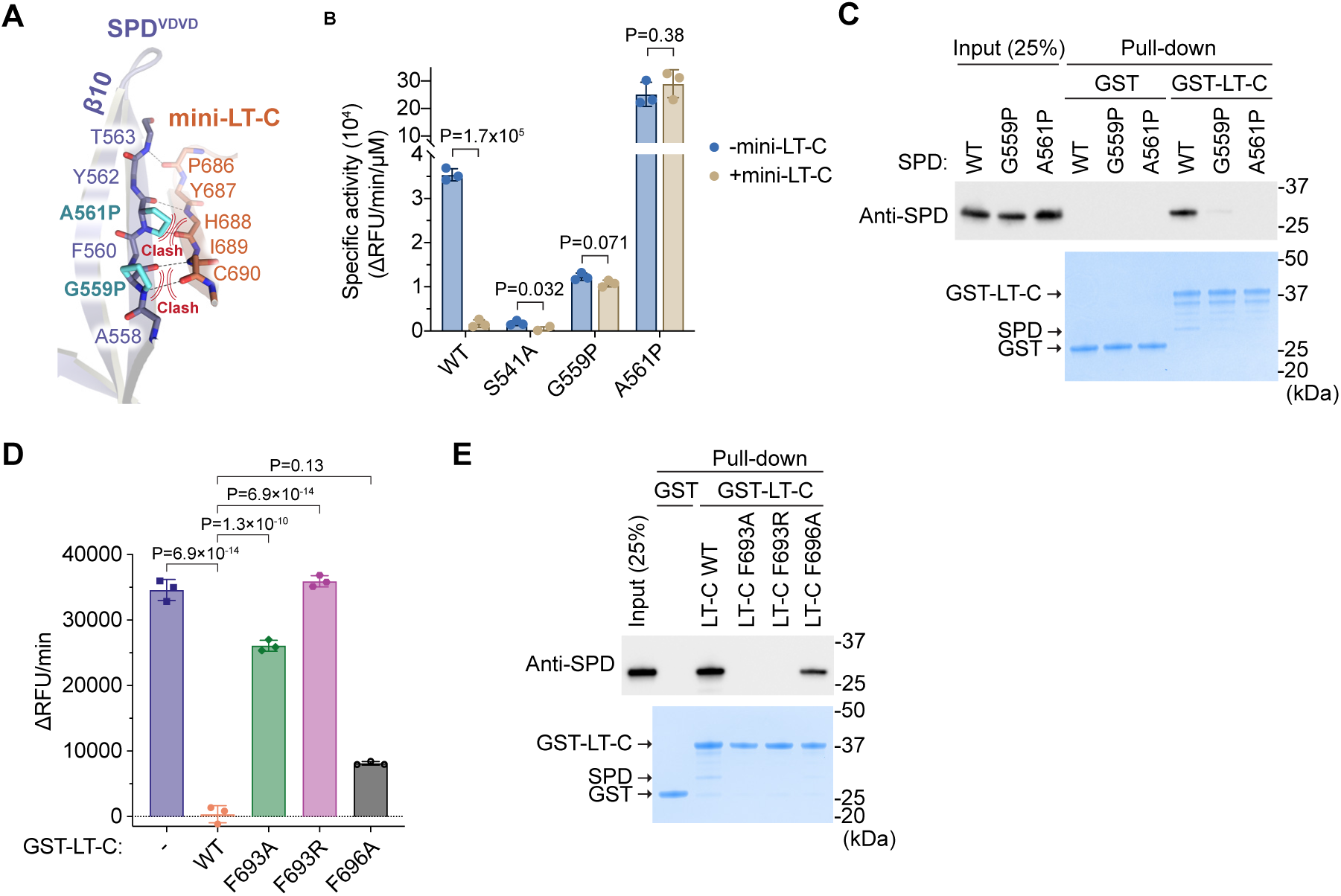
β10 interface and a P1-like residue mediate inhibition of FAM111A by SV40 LT-C. **(A)** Zoomed-in-view of the interface between SPD β10 and LT-C with G559P and A561P substitutions. Clashes at mutated residues are shown with steric hindrance indicated by curved double lines. **(B)** *In vitro* peptidase activity of FAM111A SPD (wild-type, catalytic mutant S541A, β10 mutants G559P and A561P; 1 µM) measured using Suc-AAPF-AMC as a substrate in the presence or absence of mini-LT-C (1 µM). Values are mean ± SD of three technical replicates. Statistical significance was determined by unpaired two-tailed t-test. RFU: relative fluorescence units. **(C)** *In vitro* GST pull-down assay of FAM111A SPD with GST or GST-LT-C. Bound FAM111A SPD was detected by western blotting (top), and GST proteins were visualized by Coomassie staining (bottom). Molecular weight markers are shown on the right. **(D)** *In vitro* peptidase activity of FAM111A SPD measured using Suc-AAPF-AMC as a substrate in the presence of GST-LT-C (wild-type or mutants F693A, F693R, and F696A). Reactions without GST-LT-C contain GST as a control. Values are mean ± SD of three technical replicates. Statistical significance was determined by one-way ANOVA with Dunnett’s multiple-comparison test. RFU: relative fluorescence units. **(E)** *In vitro* GST pull-down assay of FAM111A SPD with GST-LT-C (wild-type or mutants F693A, F693R, and F696A). Bound FAM111A SPD was detected by western blotting (top), and GST proteins were visualized by Coomassie staining (bottom). Molecular weight markers are shown on the right.

The crystal structure also showed that Phe693 inserts into the FAM111A S1 specificity pocket as a P1-like residue (Fig. 2A,iii). To test the functional importance of this interaction, we substituted Phe693 with alanine or arginine, while the nearby phenylalanine residue Phe696 was substituted with alanine as a control (Supplementary Fig. 6C). Based on the structure, the arginine substitution was predicted to disrupt insertion of the P1-like residue into the hydrophobic S1 specificity pocket. We then examined the effects of these mutations on FAM111A inhibition and binding. Whereas F696A caused only modest reductions in inhibitory activity, F693A markedly impaired inhibition and F693R completely abolished it (Fig. 4D and Supplementary Fig. 6D). Consistent with these results, both Phe693 substitutions disrupted the interaction between LT-C and FAM111A SPD, whereas the F696A mutant retained robust binding (Fig. 4E). These findings demonstrate that Phe693 is the principal determinant of LT-C recognition of FAM111A and support its role as a P1-like residue that engages the S1 specificity pocket.

Taken together, these findings validate the crystal structure and establish that LT-C achieves high-affinity inhibition of FAM111A through the combined engagement of an extended β-sheet interface and substrate-like recognition of the S1 specificity pocket.

### LT-C inhibits FAM111A without undergoing proteolytic cleavage

Many protease inhibitors engage their target enzymes through a substrate-like mechanism in which a P1 residue occupies the S1 pocket. Canonical (standard-mechanism) inhibitors, including members of the Kunitz, Kazal, and Bowman-Birk families, may undergo proteolytic cleavage while remaining tightly bound to the protease, thereby limiting turnover^26,27,28,29^. In contrast, serpin inhibitors, such as vaccinia virus SPI-1, undergo proteolytic cleavage as an essential step in trapping and inactivating the protease^30,31^. Given that the P1-like residue Phe693 occupies the S1 pocket of FAM111A, we next asked whether LT-C undergoes proteolytic cleavage or forms a covalent acyl-enzyme intermediate during inhibition.

Recombinant wild-type or catalytically inactive (S541A) FAM111A SPD was incubated with recombinant mini-LT-C, and the resulting complexes were purified by size exclusion chromatography (Supplementary Fig. 7A). Purified wild-type SPD remained inhibited following complex isolation (Supplementary Fig. 7B,C), allowing us to assess whether LT-C undergoes proteolysis to any detectible extent during inhibition. Liquid chromatography-mass spectrometry (LC-MS) analysis after incubation at 37°C for one hour revealed identical LT-C mass spectra in complexes containing wild-type or S541A SPD, with no evidence of LT-C cleavage (Fig. 5A). Likewise, the molecular masses of wild-type and S541A SPD remained unchanged following complex formation (Fig. 5B), indicating that LT-C neither undergoes proteolytic cleavage nor forms a covalent complex with FAM111A.

**Figure 5.**
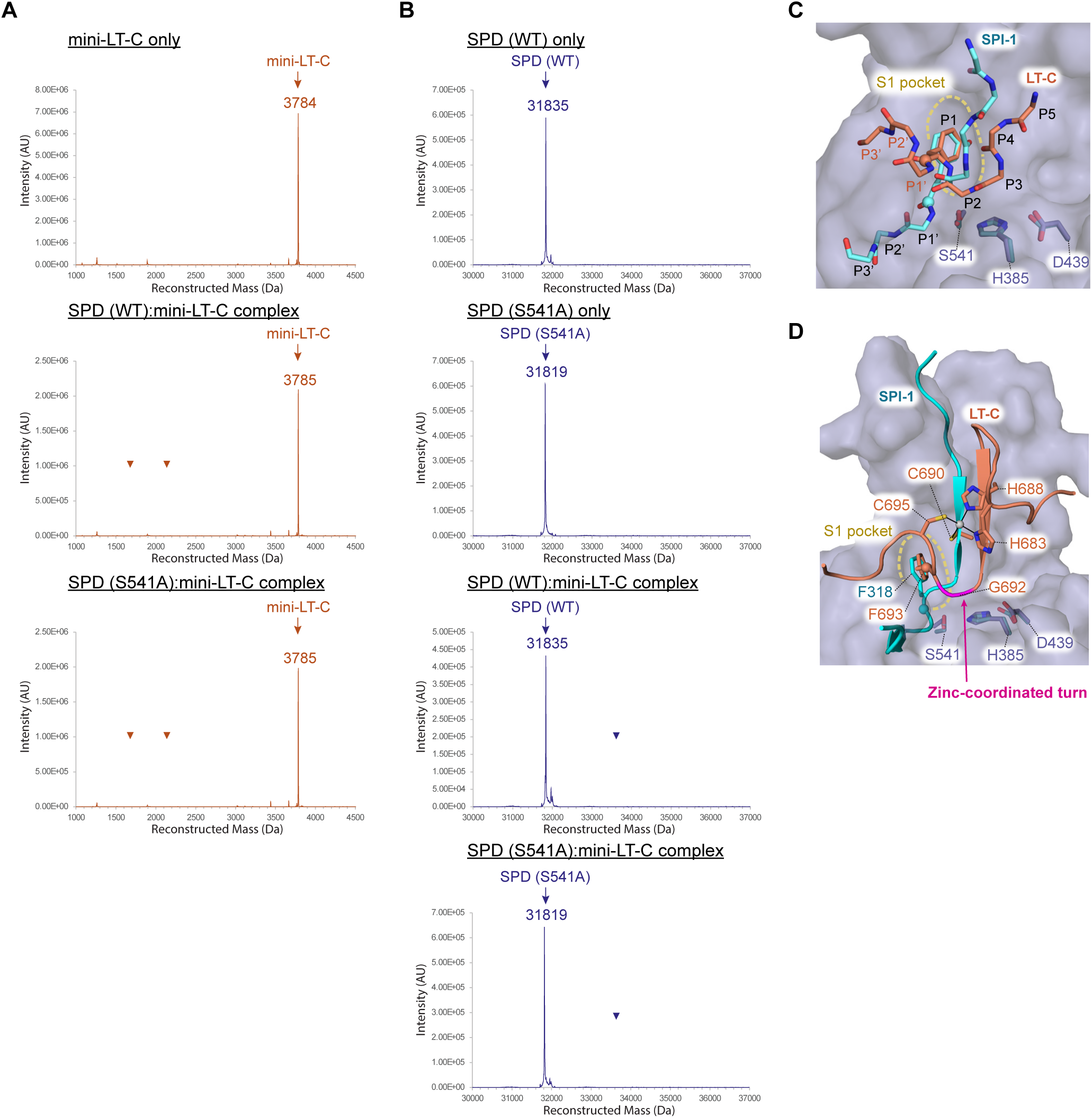
LT-C inhibits FAM111A through a cleavage-avoiding, non-covalent mechanism. **(A)** Liquid chromatography-mass spectrometry (LC-MS) analysis of mini-LT-C alone (top) and in complex with wild-type or catalytically inactive (S541A) FAM111A SPD following purification by size-exclusion chromatography. Deconvoluted mass spectra are shown. Arrowheads indicate the expected masses of N- and C-terminal cleavage products. **(B)** LC-MS analysis of wild-type or catalytically inactive (S541A) FAM111A SPD alone and in complex with mini-LT-C following purification by size-exclusion chromatography. Deconvoluted mass spectra are shown. Arrowheads indicate the expected masses of a covalent acyl-enzyme intermediate containing the N-terminal fragment of cleaved mini-LT-C. **(C)** Comparison of the binding modes of mini-LT-C (orange) and vaccinia virus serpin SPI-1 reactive center loop (cyan) within the FAM111A active site. The catalytic triad residues (H385, D439, and S541) are shown as sticks. Carbonyl carbons of the putative scissile bonds are shown as spheres. SPI-1 was modeled using a ColabFold prediction. **(D)** Structural comparison highlighting the distinct trajectories of the C-terminal region of mini-LT-C (orange) and the modeled SPI-1 reactive center loop (cyan) within the FAM111A active site. The zinc ion and coordinating residues of mini-LT-C, P1-like residue Phe693, and the catalytic triad are shown as sticks. FAM111A SPD^VDVD^ is shown as surface representation.

The crystal structure provides a structural explanation for the absence of LT-C cleavage. Although the P1-like residue Phe693 occupies the S1 pocket, the carbonyl carbon of the putative scissile bond is positioned approximately 7.4 Å from the Oγ atom of catalytic Ser541 (Fig. 5C), placing it outside the geometry required for nucleophilic attack. By contrast, ColabFold^32^-based modeling of the reactive center loop of SPI-1, a FAM111A inhibitor from vaccinia virus^19^, predicts that the P1 (Phe318) carbonyl is positioned within 2.9 Å of Ser541, consistent with the cleavage-dependent inhibitory mechanism (Fig. 5C). The zinc-coordinated turn formed around Gly692 maintains the scissile bond in a catalytically incompetent position by redirecting the C-terminal region of LT-C away from the catalytic serine (Fig. 5D). Collectively, these findings indicate that LT-C binds FAM111A as a substrate mimic while avoiding proteolytic cleavage during inhibition.

### Zinc-binding motif and Phe693 of LT-C are required for productive SV40 infection

We next asked whether the LT-C zinc-binding motif and the P1-like residue Phe693, which are required for FAM111A binding and inhibition *in vitro*, are required for the host range function of Large T during SV40 infection. To assess productive SV40 infection, we performed plaque assays using SV40 viruses encoding wild-type Large T or the C690S or F693R variants in permissive (BSC40) and non-permissive (CV-1) cells. In BSC40 cells, wild-type SV40 and both mutant viruses produced comparable numbers of plaques over a range of multiple multiplicities of infection (MOIs) (Fig. 6A,B), consistent with the established use of BSC40 cells for determining titers of host-range mutants^16,17^. By contrast, both the C690S and F693R mutant viruses exhibited a marked reduction in plaque formation in CV-1 cells (Fig. 6A,B), indicating a host range defect specifically in non-permissive cells.

**Figure 6.**
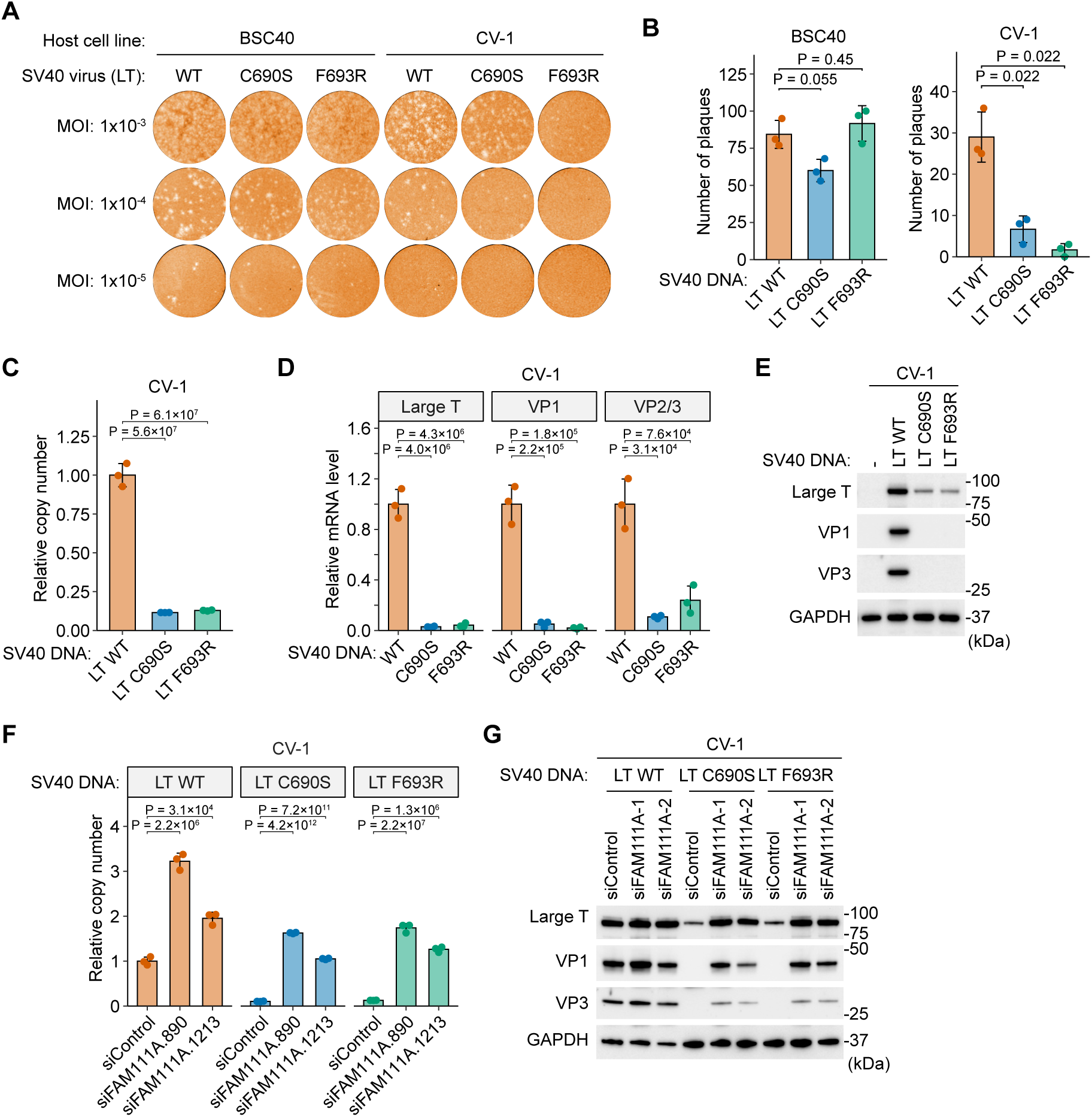
The LT-C zinc-binding motif and P1-like residue are required to overcome FAM111A-mediated host restriction. **(A)** Plaque formation assay in permissive (BSC40) and non-permissive (CV-1) cells infected with wild-type (WT) SV40 or viruses encoding Large T variants carrying the C690S and F693R mutations. Cells were infected at the indicated multiplicity of infection (MOI), and plaques were visualized 10 days post-infection and quantified at an MOI of 1×10^-4^. **(B)** Quantification of plaques from three independent experiments as described in panel A. Statistical significance was determined by one-way ANOVA with Dunnett’s multiple-comparison test. **(C)** Quantification of SV40 genome copy number in CV-1 cells transfected with WT, C690S, F693R SV40 genomic DNA. Viral genomic copy number was determined by quantitative PCR 72 hr after transfection. Values are mean ± SD of three technical replicates. Statistical significance was determined by one-way ANOVA with Dunnett’s multiple-comparison test. **(D)** Quantitative RT-PCR analysis of viral transcripts in CV-1 cells transfected with WT, C690S, or F693R SV40 genomic DNA. Relative mRNA levels of the early gene Large T and the late genes VP1 and VP2/VP3 were measured 72 h after transfection and normalized to GAPDH mRNA. Values represent mean ± SD of three independent experiments. Statistical significance was determined by one-way ANOVA with Dunnett’s multiple-comparison test. **(E)** Western blot analysis of viral protein levels in CV-1 cells transfected with WT, C690S, F693R SV40 genomic DNA. Whole-cell lysates were harvested 72 hr after transfection and analyzed using antibodies against the indicated proteins. GAPDH levels from the corresponding lysates are shown. Molecular weight markers are indicated on the right. **(F)** Quantification of SV40 genome copy number following depletion of FAM111A in CV-1 cells using two independent siRNAs (siFAM111A.890 and siFAM111A.1213). Cells were transfected with WT, C690S, or F693R SV40 genomic DNA, and viral genome copy number was determined by quantitative PCR 72 h after transfection. Values are mean ± SD of three technical replicates. Statistical significance was determined by one-way ANOVA with Dunnett’s multiple-comparison test. **(G)** Western blot analysis of viral protein levels following FAM111A depletion. Cells were treated as in (F), and whole-cell lysates harvested 72 hr after transfection were analyzed using antibodies against the indicated proteins. GAPDH levels from the corresponding lysates are shown. Molecular weight markers are indicated on the right.

To determine whether the host-range defect reflects impaired viral replication, we quantified SV40 genome amplification following transfection of equal amounts of wild-type or mutant SV40 genomic DNA into CV-1 cells. Three days after transfection, both the C690S and F693R genomes accumulated to substantially lower levels than wild type (Fig. 6C). Consistent with reduced genome amplification, expression of both the early viral gene Large T and the late genes VP1 and VP3 was markedly diminished at the mRNA and protein levels (Fig. 6D,E).

To determine whether impaired inhibition of FAM111A underlies the defects of the C690S and F693R mutants, we depleted FAM111A in CV-1 cells using two independent siRNAs (Supplementary Fig. 8A). FAM111A depletion markedly increased genome amplification of both mutants (Fig. 6F) and restored viral protein expression to levels comparable to those of wild-type SV40 (Fig. 6G). By contrast, depletion of FAM111A produced only a modest increase in wild-type genome amplification and had little effect on viral protein expression, suggesting that wild-type Large T largely overcomes FAM111A restriction. These findings demonstrate that impaired inhibition of FAM111A underlies the host-range defects of the LT-C C690S and F693R mutants during productive SV40 infection.

Together with the structural and biochemical analyses, these findings demonstrate that the zinc-binding motif and the P1-like residue Phe693 are essential for LT-C-mediated inhibition of FAM111A and establish this mechanism as a critical determinant of productive SV40 infection.

### SV40 restriction by FAM111A requires its protease activity and is partially dependent on the PIP motif

Having established the structural basis of LT-C-mediated inhibition of FAM111A, we next examined whether these interactions occur in cells. Because antibodies suitable for detecting endogenous simian FAM111A are not available, we performed these experiments in human RPE-1 cells, in which the C690S and F693R mutants exhibit genome amplification defects comparable to those observed in CV-1 cells (Supplementary Fig. 8B). Following transfection of SV40 genomic DNA, co-immunoprecipitation of Large T antigen recovered endogenous FAM111A with wild-type Large T but not with the C690S or F693R mutants (Fig. 7A). No FAM111A signal was detected in immunoprecipitates from FAM111A-knockout cells (Fig. 7A, Supplementary Fig. 8C), confirming the identity of the co-immunoprecipitated FAM111A band. These results demonstrate that both the zinc-binding motif and the P1-like residue are required for efficient LT-FAM111A interaction in cells.

**Figure 7.**
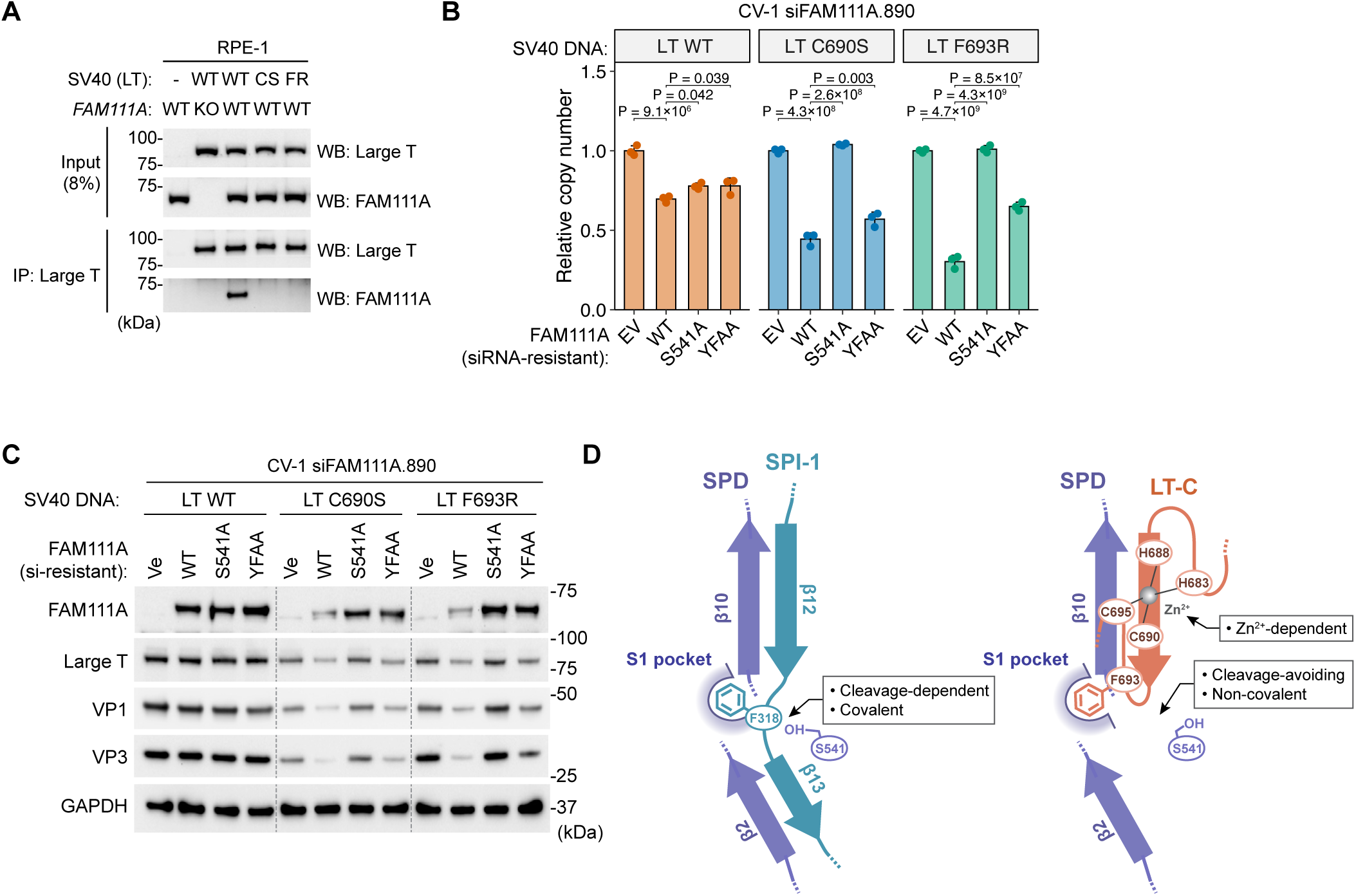
SV40 host-range restriction requires FAM111A protease activity and is partially dependent on the PIP motif. **(A)** Co-immunoprecipitation of SV40 Large T antigen and FAM111A. RPE-1 cells (FAM111A WT or KO) were transfected with SV40 genomic DNA encoding WT Large T or the C690S (CS) or F693R (FR) variants. Large T was immunoprecipitated with an anti-Large T antibody, and co-immunoprecipitated FAM111A was detected by western blotting. Input samples are shown for comparison. **(B)** Quantification of SV40 genome copy number in FAM111A-depleted CV-1 cells following cotransfection of WT, C690S, or F693R SV40 genomic DNA with empty vector (EV) or siRNA-resistant FAM111A expression plasmids encoding WT, the catalytic mutant S541A, or the PIP-box mutant YFAA. Viral genome copy number was determined by quantitative PCR 72 hr after transfection. Values are mean ± SD of three technical replicates. Statistical significance was determined by one-way ANOVA with Dunnett’s multiple-comparison test. **(C)** Western blot analysis of viral protein levels in FAM111A-depleted CV-1 cells expressing the indicated siRNA-resistant FAM111A variants following cotransfection with WT, C690S, or F693R SV40 genomic DNA. Whole-cell lysates were harvested 72 hr after transfection and analyzed using antibodies against the indicated proteins. GAPDH levels from the corresponding lysates are shown. Molecular weight markers are indicated on the right. **(D)** Model for FAM111A inhibition by the poxvirus serpin SPI-1 and SV40 Large T antigen C-terminal domain (LT-C). Both SPI-1 (teal) and LT-C (orange) inhibit the FAM111A SPD (purple) through a common recognition mechanism involving formation of an antiparallel β-sheet and insertion of a P1 residue into the S1 specificity pocket. SPI-1 functions as a classical suicide substrate, in which the reactive center loop is cleaved by FAM111A, resulting in formation of a covalent inhibitory complex. In contrast, LT-C employs a Zn^2+^-dependent structural motif that positions the P1-like residue Phe693 within the S1 specificity pocket while preventing catalytic cleavage, thereby forming a stable non-covalent inhibitory complex.

We next asked whether FAM111A-mediated restriction requires its protease activity and PCNA interaction. To address this question, endogenous FAM111A was depleted using siFAM111A.890 and replaced with siRNA-resistant wild-type FAM111A, the catalytic mutant S541A, or the PIP-box mutant YFAA. For SV40 genomes encoding the LT C690S or F693R mutation, restriction was restored by re-expression of wild-type FAM111A but not by the catalytic mutant S541A, indicating that FAM111A-mediated restriction of these LT mutants requires its protease activity (Fig. 7B). By contrast, the PIP-box mutant YFAA only partially restored restriction, resulting in intermediate levels of viral DNA replication, indicating that PCNA interaction contributes to, but is not absolutely required for, FAM111A-mediated restriction. Consistent with these findings, expression of viral proteins from SV40 genomes encoding the LT C690S or F693R mutations was strongly suppressed by wild-type FAM111A, unaffected by the catalytic mutant S541A, and only partially suppressed by the PIP-box mutant YFAA (Fig. 7C). Notably, wild-type FAM111A protein levels were consistently reduced in cells transfected with the LT C690S or F693R mutant genomes, whereas the catalytically inactive S541A mutant remained stable, suggesting that FAM111A protease activity may contribute to the reduced abundance of wild-type FAM111A under these conditions. These findings demonstrate that FAM111A restricts SV40 through its protease activity and that efficient restriction is enhanced by its PCNA interaction.

## Discussion

In this study, we identify the C-terminal region of SV40 Large T antigen as a previously unrecognized inhibitor of the host serine protease FAM111A and define the structural mechanism by which it overcomes FAM111A-mediated host restriction. By combining structural, biochemical, and virological approaches, we show that LT-C engages the FAM111A active site through a zinc-dependent, cleavage-avoiding mechanism that requires an H2C2-like zinc-binding motif and a P1-like phenylalanine residue. Disruption of either structural feature abolishes FAM111A binding and inhibition, resulting in impaired viral genome amplification, reduced viral gene expression, and host-range defects in non-permissive cells. Conversely, depletion of FAM111A rescues these defects, whereas re-expression of wild-type FAM111A, but not the catalytically inactive mutant, restores viral restriction. Altogether, these findings establish direct inhibition of the FAM111A protease as a key mechanism by which SV40 overcomes host restriction.

LT-C inhibits FAM111A through a mechanism that shares features with substrate-mimicking protease inhibitors but is mechanistically distinct. Like the poxvirus serpin SPI-1, LT-C engages FAM111A through substrate-like interactions in which a P1-like residue occupies the S1 specificity pocket. However, whereas SPI-1 undergoes proteolytic cleavage followed by covalent trapping of FAM111A (Fig. 7D, left), LT-C avoids proteolysis altogether (Fig. 7D, right). More broadly, this also distinguishes LT-C from canonical (standard-mechanism) inhibitors, including members of the Kunitz, Kazal, and Bowman-Birk families, which remain inhibitory despite proteolytic cleavage because the cleaved inhibitor remains tightly bound, allowing religation of the scissile bond. Our crystal structure shows that, although the P1-like phenylalanine of LT-C occupies the FAM111A S1 pocket, the putative scissile bond is displaced from the catalytic serine into a catalytically incompetent configuration. Thus, rather than relying on or tolerating proteolytic cleavage, LT-C forms a stable inhibitory complex by uncoupling substrate recognition from catalysis.

Another striking feature of LT-C-mediated inhibition is its dependence on zinc coordination. LT-C contains a H2C2-like zinc-binding motif that is essential for FAM111A inhibition, and disruption of this motif abolishes the host-range function of Large T. Although zinc-binding motifs are widely used by viral proteins to mediate protein-protein interactions^33^, the LT-C motif instead plays a direct structural role in protease inhibition. Specifically, zinc coordination stabilizes a turn within the C-terminal region of LT-C that redirects the reactive loop away from the catalytic serine while preserving engagement of the P1-like residue with the S1 pocket. This geometry enables LT-C to bind the FAM111A active site as a substrate mimic while maintaining the putative scissile bond in a catalytically incompetent configuration. To our knowledge, this represents a previously unrecognized mechanism of protease inhibition in which metal coordination stabilizes a cleavage-avoiding conformation of an active-site-bound inhibitor.

FAM111A has previously been implicated in DNA replication-associated processes, including protection of replication forks from protein-DNA obstacles^22,24^. Our data demonstrate that FAM111A-mediated restriction of SV40 requires its protease activity and is partially dependent on its PCNA-interacting PIP motif, indicating that efficient antiviral activity is facilitated by, but does not strictly require, PCNA binding. Consistent with this observation, FAM111A-mediated restriction of vaccinia virus also depends on its protease activity, although PCNA interaction was reported to be dispensable in that system^34^. Together, these findings suggest that FAM111A antiviral activity is fundamentally driven by its protease function but may be enhanced by replication fork-associated localization in certain contexts. Previous studies reported a modest reduction in viral DNA replication in LT-C host-range mutants but concluded that the primary defect occurred at later stages of infection^13,14,15,35,36^. Our findings instead suggest that inhibition of viral genome amplification by FAM111A represents an earlier and more significant component of the host-range phenotype than previously appreciated. An additional intriguing observation is that the abundance of ectopically expressed wild-type FAM111A was consistently reduced in cells transfected with the LT-C C690S and F693R mutant genomes, whereas the catalytically inactive S541A mutant remained stable. Although the underlying mechanism remains to be determined, this observation raises the possibility that productive engagement of FAM111A promotes protease-dependent turnover, potentially through autocleavage or another activation-associated mechanism. Future studies will be required to determine the physiological substrates of FAM111A during SV40 infection and how its protease activity is coordinated with viral DNA replication. Additionally, because Large T antigen is widely used to immortalize and transform cells, its ability to inhibit FAM111A may also influence host DNA replication dynamics and DNA repair processes, raising the possibility that FAM111A inhibition contributes to the cellular effects of Large T beyond viral infection.

Not all polyomaviruses encode a C-terminal host-range domain in Large T antigen, indicating that this region is not universally required for viral replication but instead represents an adaptive feature acquired by a subset of viruses. Building on the previous identification of FAM111A as a host-range restriction factor antagonized by the Large T C-terminus^1^, the present study defines the molecular basis of this antagonism by showing that LT-C directly inhibits the FAM111A protease through a zinc-dependent mechanism. The resulting defects in viral genome amplification, gene expression, and productive infection caused by disruption of this inhibitory interface, together with their rescue by FAM111A depletion, establish direct protease inhibition as the mechanism by which LT-C overcomes FAM111A-mediated restriction. The selective retention of the Large T C-terminal domain in SV40, JCV, BKV, and SA12^17^ raises the possibility that divergence of LT-C sequences among polyomaviruses reflects adaptation to distinct host restriction environments. Our findings support a model in which viral host range is shaped by the evolution of specific antagonists that neutralize individual host restriction factors, exemplified here by inhibition of the FAM111A protease.

In summary, our study defines a zinc-dependent, cleavage-avoiding mechanism by which SV40 Large T antigen inhibits the host restriction factor FAM111A. These findings provide a mechanistic explanation for the long-standing host-range function of the Large T C-terminus. Our work further suggests that host serine proteases involved in DNA replication can also function as antiviral restriction factors and that viral inhibition of these enzymes represents an effective strategy for expanding host range. The independent evolution of mechanistically distinct FAM111A inhibitors in polyomaviruses and poxviruses^1,2^, represented by the cleavage-avoiding LT-C inhibitor described here and the cleavage-dependent serpin SPI-1^19^, highlights the importance of FAM111A as an antiviral defense factor. Collectively, these findings extend our understanding of how DNA viruses adapt to restrictive host environments and identify protease inhibition as a potentially widespread mechanism of viral counter-defense.

## Methods

### Plasmids and SV40 DNA

For bacterial expression, a codon-optimized human FAM111A SPD sequence was cloned into pDB.His.MBP as described previously^22^. DNA fragments encoding SV40 LT amino acids 627-708 (LT-C) and 654-708 were amplified by PCR and cloned into pGEX-6P-2. DNA fragments encoding SV40 LT amino acids 665-708, 674-708, and 679-707 (mini-LT-C) were synthesized (IDT; Supplementary Table 2) and cloned into pGEX-6P-2. SV40 genomic DNA was a gift from Dr. Barry Milavetz (University of North Dakota) and cloned into the pUC18 vector. Point mutations were introduced into constructs using QuikChange (Agilent). Plasmid sequences were confirmed by Sanger or whole plasmid sequencing. pMD2.G and psPAX2 were gifts from Didier Trono. Primers and plasmids used in this study are listed in Supplementary Tables 3 and 4, respectively.

### Bacterial cell culture

For bacterial expression of His_6_.MBP-FAM111A SPD, *E. coli* BL21(DE3) cells harboring pDB.His.MBP/FAM111A SPD were grown in Terrific Broth at 37°C in a Lex-10 or Lex-48 bioreactor (Epiphyte3). When cultures reached an optical density (OD_600_) of 1.2-1.5, protein expression was induced with 50 µM isopropyl b-D-1-thiogalactopyranoside (IPTG) supplemented with 2% (v/v) glycerol and 1% (v/v) ethanol, followed by incubation at 10°C for approximately 40 hr. For bacterial expression of GST-LT-C proteins, *E. coli* BL21(DE3) carrying pGEX-6P-2/LT-C were grown in Terrific Broth at 37°C in an orbital shaker until cultures reached at OD_600_ of 0.6-0.8. Expression of GST-LT-C was induced by the addition of 200 µM isopropyl b-D-1-thiogalactopyranoside (IPTG), and cells were cultured at 18°C for 21 hours. Bacterial cells were harvested by centrifugation (3,200 × g, 25 min, 4°C), and cell pellets were stored at −80°C until protein purification.

### Protein purification

FAM111A SPD proteins were purified as described previously^19,22^. Briefly, bacterial cell pellets were resuspended in Ni-NTA binding buffer (50 mM sodium phosphate pH 7.5, 500 mM NaCl, 5 mM imidazole, 0.5 mM Tris(2-carboxyethyl)phosphine hydrochloride (TCEP), 5% glycerol, 250 µg/ml lysozyme), lysed by sonication and clarified by centrifugation. The clarified lysate was purified using Ni-NTA resin followed by amylose resin. Eluted protein was precipitated by adding two volumes of 4 M ammonium sulfate overnight on ice. The precipitated protein was collected by centrifugation, resuspended, and further purified by size exclusion chromatography using a HiLoad 16/600 Superdex 200 pg column (S200) (Cytiva) pre-equilibrated with gel filtration buffer (20 mM Tris pH 7.4, 300mM NaCl, and 1 mM TCEP). Fractions containing the protein were pooled, and the His_6_-MBP tag was cleaved off with TEV protease for 2-3 days on ice. The cleaved FAM111A SPD was separated from the tag by anion exchange chromatography using a RESOURCE Q (ResQ) column (Cytiva). Purified protein was aliquoted, flash frozen in liquid nitrogen, and stored at −80°C until use.

For FAM111A SPD used in the streptavidin pull-down assays, the same purification procedure was followed with the inclusion of EDTA to minimize zinc contamination. Specifically, 1 mM EDTA was added to buffers used for amylose affinity and S200 size exclusion chromatography, and 0.5 mM EDTA was included to ResQ buffers. Following ResQ purification, the protein was buffer exchanged to gel filtration buffer supplemented with 25% glycerol and concentrated using Amicon Ultra centrifugal filters (Millipore).

GST-LT-C proteins used for pull-down assays were purified using Glutathione Sepharose 4B beads (Cytiva) by batch purification. Bacterial cell pellets were resuspended in PBS containing 250 μg/ml lysozyme and lysed by sonication. Lysates were clarified by centrifugation (17,200 x g, 45 min, 4°C) and incubated with glutathione beads for 2 hr at 4°C. Beads were washed four times with wash buffer (50 mM Tris-HCl pH 7.5, 250 mM NaCl, 10% glycerol) then eluted with the wash buffer supplemented with 20 mM glutathione. Glutathione was removed using Zeba spin 7K MWCO desalting columns (Thermo Fisher Scientific), and 1 mM TCEP was added to the final buffer. Proteins were aliquoted and flash frozen in liquid nitrogen until use.

GST and GST-LT-C proteins used for inductively coupled plasma mass spectrometry (ICP-MS) underwent a more extensive purification scheme using an ÄKTA Pure system. After clearing lysate by centrifugation, samples were loaded onto a GSTrap affinity column (Cytiva). Unbound proteins were washed out with lysis buffer, followed by GSTrap wash buffer (50 mM Tris pH 7.4, 500 mM NaCl, 10% glycerol). Purified proteins were eluted with GSTrap elution buffer (50 mM Tris pH 7.4, 300 mM NaCl, 10% glycerol, 20 mM glutathione, buffer pH adjusted to 7.0). Eluted fractions were pooled and further purified by size-exclusion chromatography on an S200 column.

To produce LT-C peptides, peak fractions from a GSTrap column containing GST-mini-LT-C peptide (amino acids 679-708) were pooled and were further purified by S200 pre-equilibrated with gel filtration buffer. The GST tag was removed by overnight cleavage with 3C protease (Genscript) on ice, and the cleaved mini-LT-C peptide was separated from the GST tag and protease by a second size-exclusion step using a Superdex 75 pg column (Cytiva) equilibrated with gel filtration buffer. Fractions containing purified mini-LT-C peptide were pooled, concentrated, aliquoted, flash frozen in liquid nitrogen, and stored at −80°C until further use.

SPD:mini-LT-C complexes were prepared by mixing purified SPD (wild-type, S541, or VDVD) and mini-LT-C proteins at a molar ratio of 1:1.5 and incubating the mixture overnight at 4°C. The resulting complex was separated from unbound SPD protein and mini-LT-C peptides by size-exclusion chromatography using a Superdex 200 Increase 10/300 analytical column (Cytiva) pre-equilibrated with gel filtration buffer containing 20 mM Tris pH 8.0, 150 mM NaCl, 1 mM DTT. Fractions corresponding to the SPD:mini-LT-C complex were pooled, concentrated, aliquoted, flash frozen in liquid nitrogen, and stored at −80°C until further use.

Protein concentrations were measured using a Qubit 4 fluorometer with Qubit Protein Assay Kit (Thermo Fisher Scientific), Bradford reagent (Bio-Rad), or absorbance at 280 nm using calculated extinction coefficients. One microgram of each purified recombinant protein was analyzed by SDS-PAGE to assess the purity and concentration. All protein gels were stained with AcquaStain protein gel stain (Bulldog Bio), imaged with ChemiDoc (Bio-Rad), and analyzed with Image Lab software (Bio-Rad, version 6.1.0).

### *In vitro* peptidase assay

Peptidase assays were performed using an AMC-conjugated substrate (Suc-AAPF-AMC; Sigma #230914-25MG) in black 384-well plates. All reactions were carried out in a total volume of 25 µL and contained 0.2 mg/ml BSA, 1 µM FAM111A SPD, 2.5 µM GST-LT-C protein, and 1 mM substrate unless otherwise stated. For protease assays using synthetic core-LT-C peptides, peptides corresponding to residues 679-704 were used at a final concentration of 5 µM together with 2.5 µM ZnCl_2_. The core-LT-C peptide sequences used are listed in Supplementary Table 5. For all reactions, FAM111A SPD was pre-incubated with the indicated amount of GST-LT-C protein or LT-C peptides for 10 min at 37°C unless otherwise stated. Reactions were initiated by addition of substrate, and fluorescence (Ex/Em: 380 nm/460 nm) was measured every two minutes for 1 hour at 37°C using the SpectraMax i3x plate reader (Molecular Devices).

### *In vitro* pull-down assays

For the *in vitro* GST pull-down assays, 5 µg of GST tagged proteins were incubated with 15 µL Glutathione Sepharose 4B beads (bed volume; Cytiva) for 30 min at 4°C in pull-down buffer (20 mM Tris-HCl (pH 7.4), 150 mM NaCl, 1% NP40, 1 mM DTT, 25 µg/ml BSA). Beads were washed once and then incubated for 1 hr at 4°C with 1 µg of FAM111A SPD protein in 800 µL pull-down buffer. Subsequently, beads were washed five times with pull-down buffer and bound proteins were eluted by boiling with 2x SDS sample buffer. Pull-down samples were visualized using AcquaStain protein gel stain (Bulldog Bio) after SDS-PAGE to detect GST-tagged proteins, or by western blotting using an anti-FAM111A SPD antibody (custom-made, Cocalico) to detect FAM111A SPD^19^.

*In vitro* streptavidin pull-down assays were performed using streptavidin agarose beads (Millipore) and biotinylated LT-C peptides corresponding to residues 679-704 (GenScript). The core-LT-C peptide sequences used are listed in Supplementary Table 5. Where indicated, peptides were pre-incubated with 2-fold molar excess of ZnCl_2_ for 30 min prior to binding to the streptavidin beads. For assays performed in the presence of EDTA, 0.5 mM EDTA was included in the pull-down buffer.

### Fluorescence polarization anisotropy

Fluorescein-labelled core-LT-C (FITC-core-LT-C) synthetic peptide was dissolved in water and added to a folding reaction consisting of 10 mM Tris pH 7.5, 0.5 mM TCEP, and 20 μM ZnCl_2_, with a final FITC-core-LT-C peptide concentration of 10 μM, and incubated overnight at 4°C. The folded FITC-core-LT-C peptide was then purified by anion exchange chromatography, by binding it to a ResQ 6 mL column (Cytiva) in low salt buffer (20 mM Tris pH 7.5, 3 mM DTT) with a gradient from 0 to 60% high salt buffer (20 mM Tris pH 7.5, 1M NaCl). FITC-core-LT-C eluted as a single peak around 150 mM NaCl, and was concentrated and buffer exchanged into storage buffer (20 mM Tris pH 8.0, 270 mM NaCl, 0.5 mM TCEP, and 25% v/v glycerol) using a 3 kDa molecular weight cutoff centrifugal concentrator (Millipore). SPD proteins used for FP analysis were similarly exchanged into the same storage buffer, but with a 10 kDa molecular weight cutoff centrifugal concentrator (Millipore).

Fluorescent polarization anisotropy (FP) assays were performed essentially as described^37^. Briefly, FITC-core-LT-C peptide (1 nM) was incubated with the indicated concentrations of SPD proteins in triplicate (N = 3) in FP buffer (20 mM Tris pH 8, 270 mM NaCl, 0.5 mM TCEP, 0.2 mg/mL BSA and 5% (v/v) glycerol) in black, flat-bottomed 96-well plates for 2 hours at 22 °C, then each replicate was measured eight times in a Clariostar+ microplate reader (BMG Labtech) using excitation and emission wavelengths of 485 and 520 nm, respectively. To measure FP of LT-C peptide in the absence of zinc, the same assay was performed but without pre-binding zinc to the LT peptide, and the inclusion of 1 mM EDTA in the FP binding reaction. K_D_ values were calculated by fitting FP data to a one-site binding model in GraphPad Prism.

### Protein crystallization, data collection, and structure determination

SPD^VDVD^ protein was concentrated to 17 mg/mL after Q-column purification using a 10 kDa molecular weight cutoff centrifugal concentrator (Millipore). The co-purified FAM111A SPD^VDVD^:mini-LT-C complex was concentrated to 10 mg/mL using a 3 kDa molecular weight cutoff centrifugal concentrator (Millipore). Initial crystallization screening was performed using commercially available sparse-matrix screens (Jena Bioscience, Hampton Research, and Molecular Dimensions) by the sitting-drop vapor diffusion method at 20°C. Crystals of SPD^VDVD^ were grown in 100 mM HEPES pH 7, 200 mM MgCl2, and 16-20% (w/v) PEG6000 at 4°C in drops of 200 nL protein and 200 nL reservoir solution. For the SPD^VDVD^:mini-LT-C complex, drops consisting of 200 nL protein solution and 200 nL reservoir solution were dispensed at protein-to-reservoir ratios of 1:1, 1:2, and 2:1. Initial crystallization hits produced thin needle-like microcrystals under several conditions at the 1:1 and 2:1 mixing ratios. These conditions were subsequently optimized by varying precipitant composition and increasing the protein concentration to improve crystal morphology and diffraction quality. Diffraction-quality crystals of the SPD^VDVD^:mini-LT-C complex were obtained using protein concentrated to 17.5 mg/mL in a reservoir solution containing 100 mM BIS-TRIS (pH 5.5), 200 mM MgCl₂, and 25% (w/v) polyethylene glycol (PEG) 3350. Crystals were cryoprotected by brief transfer into reservoir solution supplemented with 25% (w/v) glycerol, flash-cooled in liquid nitrogen, and transported to the NE-CAT beamlines at the Advanced Photon Source (APS), Argonne National Laboratory, for X-ray diffraction data collection. Diffraction data were collected at beamline 24-ID-E using the beamline control software and processed and scaled with HKL2000 (version 7.2.0).

The structures were solved by molecular replacement using PHENIX (version 1.17.1-3660), with the crystal structure of chain D from S541A SPD (PDB entry 8S9K) used to solve the SPD^VDVD^ apo structure, and the SPD^VDVD^ apo protein was subsequently used as the search model for the SPD^VDVD^:mini-LT-C complex structure. Molecular replacement yielded single unambiguous solutions with a translation function Z-score (TFZ) of 38.3 and 87.5 and log-likelihood gain (LLG) values of 2,347 and 10,931 respectively for the two structures. The model was iteratively improved through manual rebuilding in Coot (version 0.8.9.1) followed by refinement in PHENIX.REFINE against the full-resolution diffraction data. Data collection and refinement statistics are summarized in Supplementary Table 1. Structural figures were prepared using PyMOL (version 3.1.0, Schrödinger) and UCSF ChimeraX (version 1.10). Structural alignments, root-mean-square deviation (RMSD) calculations were performed in PyMOL and ChimeraX.

### Inductively Coupled Plasma Mass Spectrometry (ICP-MS)

Wet, acid-based digestion was performed on GST and GST-LT-C recombinant proteins. Briefly, 50 µL of buffer, GST protein, or GST-LT-C proteins were combined with 100 µL trace metal grade nitric acid (EMD Millipore) and heated to 90°C for two hours. Following, 50 µL of trace metal grade hydrogen peroxide (Sigma Aldrich) was added to each tube and heated for an additional hour. Upon cooling to room temperature, 800 µL of Milli-Q water, spiked with elemental yttrium (Inorganic Ventures) was added and the sample vortexed. Samples were then injected via peristaltic pump into a Thermo iCAP Q ICP-MS (Thermo Fisher Scientific) where data was collected in KED mode, specifying quantitation of Zn^67^ and Y^89^. A two segment, 2-fold standard curve was constructed using elemental zinc standards (Inorganic Ventures) with spiked-in elemental yttrium, allowing for quantitation. Analysis was performed across three distinct days, each time in technical triplicate, for a total of 9 data points per protein of interest. Baseline correction was performed by subtracting out average zinc present in the buffer solution by analysis day.

### Liquid Chromatography-Mass Spectrometry (LC-MS)

Samples were diluted 10-fold 30% acetonitrile and 0.05% trifluoroacetic acid to denature both protein and peptide. Masses were then measured on an X500B quadrupole time-of-flight mass spectrometer (SCIEX) coupled to an Exion liquid chromatography system (SCIEX). Peptides and proteins were separated on Aeris Widepore XB-C8 column (Phenomenex) at a flow rate of 300 µl using the following gradient from 0.1% formic acid in water (Buffer A) to 0.1% formic acid in acetonitrile (Buffer B): 2% B to 40% B over 4.5 min, 40% B to 90% B over 0.5 min. Protein and peptide were eluted directly into the mass spectrometer and mass spectra collected in positive ion mode over the range 400-2000 Da. The source parameters were: source gas 1 – 35 psi; source gas 2 – 35 psi; curtain gas – 30 psi; CAD gas 7 psi; temperature 300 °C; spray voltage 5200 V; declustering potential 120 V. Mass reconstruction was performed using the Bio Tool Kit functionality of Explorer in SCIEX OS (SCIEX).

### Mammalian cell culture

African green monkey cell lines BS-C-1 and CV-1 were cultured in Eagle’s Minimum Essential Medium (EMEM) supplemented with 1 mM sodium pyruvate. BSC40, a high-temperature tolerant derivative of BS-C-1^38^ was cultured in Dulbecco’s Modified Eagle Medium (DMEM). The human retinal pigment epithelial cell line RPE-1 was cultured in DMEM/F12 supplemented with 0.01 mg/ml hygromycin B. All media were supplemented with 10% fetal bovine serum (FBS). BS-C-1 was a gift from Dr. Bernard Moss, and BSC40 (#CRL-2761), CV-1 (#CCL-70), and RPE-1 (#CRL-4000) were purchased from American Type Culture Collection. All cell lines were confirmed to be mycoplasma-negative using the LookOut *Mycoplasma* PCR Detection Kit (Sigma).

Transfection of SV40 genomic DNA was performed using Lipofectamine 2000 (Thermo Fisher Scientific). siRNA transfections were carried out by reverse transfection using RNAiMAX (Thermo Fisher Scientific). Target sequences of siRNAs are listed in Supplementary Table 3.

### Generation of knockout cell lines

FAM111A knockout RPE-1 cell lines were generated by transfection of ribonucleoprotein (RNP) complexes containing Cas9 nickase (NEB) and paired sgRNAs targeting exon 4 of FAM111A (Supplementary Table 3) using the Nucleofector 4D system (Lonza) according to the manufacturer’s instructions. RNP complexes were assembled using 40 pmol Cas9 nickase and 60 pmol of each sgRNA (120 pmol sgRNA, Supplementary Table 3) prior to transfection. Following nucleofection, single-cell clones were isolated and expanded. Loss of FAM111A expression was confirmed by immunoblotting using an anti-FAM111A antibody. Disruption of the FAM111A locus was further verified by PCR amplification of genomic DNA surrounding the sgRNA target sites, followed by sequencing of the PCR amplicons. The primers used for genotyping are listed in Supplementary Table 3.

### SV40 infection

To generate SV40 viral stocks, wild-type and mutant SV40 genomes were excised from the pUC18 vector by restriction enzyme digestion, re-circularized with T4 ligase (NEB), and transfected into BS-C-1 cells (30 ng SV40 DNA per transfection). Cells were harvested at 12 days post transfection, when strong signs of cytopathic effect were observed. To release virus, cell pellets were subjected to three freeze-thaw cycles (ethanol/dry ice bath and 37°C water bath), followed by sonication for two cycles at 30% amplitude (30 sec on, 30 sec off). Viral stocks were aliquoted and stored at −80°C until use.

For plaque assays, BSC40 and CV-1 cells were seeded in 6-well plates and infected with SV40 viral stocks at specified MOIs, which were serially diluted in EMEM supplemented with 1 mM sodium pyruvate and 2% FBS (EMEM-2). After 2 hr, the infection medium was removed, and cells were overlaid with a 1:1 mixture of 2xEMEM without phenol red (supplemented with 2 mM sodium pyruvate, 4 mM L-glutamine and 10% FBS) and 1.8% Bacto Agar (BD) equilibrated to 42°C. Every 3 days, an additional layer of the 2×EMEM/agar mixture was added on top of the existing overlay. At 6 days post infection (dpi), neutral red was added to the agar mixture to a final concentration of 50 µg/ml. Plaques were imaged at 10 dpi using the ChemiDoc MP (Bio-Rad). SV40 wild-type and mutant viral titers were determined by plaque assays using the permissive BSC40 cell line.

### SV40 genome copy number quantification by qPCR

SV40 genomic DNA was isolated from cells transfected with SV40 genomic DNA using a modified Hirt extraction method^39,40^. Briefly, cells harvested from a 12-well plate were resuspended in 250 µL of Buffer I (50 mM Tris-HCl, pH 7.5; 10 mM EDTA; 50 µg/mL RNase A; 0.1% RNase T1 cocktail (Ambion). 250 µL of Buffer II (1.2% SDS) was then added, and samples were incubated for 5 min at room temperature. Subsequently, 350 µL of Buffer III (3 M CsCl, 1 M potassium acetate, 0.67 M acetic acid) was added, followed by incubation for 10 min at room temperature. Samples were centrifuged at 16,000 × g for 10 min at room temperature, the tubes were inverted several times, and centrifuged again at 16,000 × g for an additional 10 min. The supernatant was loaded onto Qiagen Miniprep columns (Qiagen), and columns were washed with 750 µL of wash buffer (60% ethanol, 10 mM Tris-HCl pH 7.5, 50 µM EDTA, 80 mM potassium acetate). Columns were centrifuged at 16,000 × g for 5 min at room temperature to remove residual wash buffer, and DNA was eluted in 75 µL of elution buffer (10% Qiagen EB buffer, 0.1 mM EDTA). To remove linear fragments of cellular DNA, 40 µL of the eluted DNA was treated with 4 U of DpnI (NEB) and 3 U of ATP-dependent DNase (Exonuclease V; NEB) in a 50 µL reaction containing 1 mM ATP. Reactions were incubated for 1 h at 37°C, followed by heat inactivation at 70°C for 30 min.

SV40 genome copy numbers were determined by qPCR using previously published primers targeting the SV40 genome in the VP2/VP3 genes (Supplementary Table 3)^41^. qPCR reactions were performed in technical triplicate in a total volume of 20 µL, containing 1 µM of each primer, 10 µl of PowerTrack SYBR Green Master Mix (Thermo Fisher Scientific), and 5 µL of DNA extract (5 % of total recovered DNA). Reactions were run using a QuantStudio 5 Real-Time PCR System (Thermo Fisher Scientific) using the following amplification conditions: 2 min at 50°C, 10 min at 95°C, followed by 40 cycles of denaturation at 95°C for 15 sec and annealing/extension at 60°C for 1 min. A standard curve was generated using the pUC18/SV40 wild-type plasmid as a template, with known SV40 copies ranging from 10^2^-10^10^ copies per reaction, to calculate SV40 copy numbers from unknown samples.

### RNA isolation and quantitative RT-PCR (qRT-PCR)

Total RNA was isolated from cells using TRIzol (Thermo Fisher Scientific) according to the manufacturer’s instructions. Briefly, cells were lysed directly in TRIzol, followed by phase separation with chloroform, RNA precipitation with isopropanol, washing with 75% ethanol, and resuspension in RNase-free water. RNA concentration and purity were determined by spectrophotometry.

For reverse transcription, 1 µg of total RNA was converted to cDNA using SuperScript IV VILO Master Mix (Thermo Fisher Scientific) according to the manufacturer’s protocol. Quantitative PCR was performed using PowerTrack SYBR Green Master Mix (Thermo Fisher Scientific) on a QuantStudio 5 Real-Time PCR System (Thermo Fisher Scientific) in technical triplicate using gene-specific primers (Supplementary Table 3). The following amplification condition was used: 2 min at 50°C, 10 min at 95°C, followed by 40 cycles of denaturation at 95°C for 15 sec and annealing/extension at 60°C for 1 min. Relative mRNA levels were calculated using the ΔΔCt method and normalized to GAPDH expression.

### Western blotting and co-immunoprecipitation

SV40 infected cells and SV40 DNA transfected cells were lysed directly in 12-well plates using 2xSDS sample buffer, and 5% of the harvested lysates were used for western blotting. Proteins were separated by SDS-PAGE and transferred to nitrocellulose membranes. Membranes were blocked with 5% nonfat dry milk in PBST and probed with indicated primary antibodies. After incubation with appropriate secondary antibodies, signals were detected by chemiluminescence, imaged using the ChemiDoc MP (Bio-Rad), and analyzed using Image Lab software (Bio-Rad, version 6.1.0). Primary antibodies used for western blotting are listed in Supplementary Table 6.

For coimmunoprecipitation, cells were harvested and lysed in NP-40 lysis buffer (50 mM Tris-HCl pH 7.4, 150 mM NaCl, 0.5% NP-40, 10% glycerol, 5 mM EDTA, 50 mM NaF, 1 mM Na₃VO₄) supplemented with a protease inhibitor cocktail (Sigma). Lysates were treated with 250 U/mL Benzonase (Sigma) and sonicated and clarified by centrifugation. Cell lysates containing 1 mg of total protein were incubated overnight at 4°C with rotation with 1 µg of mouse anti-SV40 Large T antigen antibody (Supplementary Table 6). Immune complexes were captured by incubation with Protein G agarose (Cytiva) for 2 hr at 4°C with rotation. Beads were washed five times with NP-40 lysis buffer, and bound proteins were eluted in 2×SDS sample buffer containing 4% β-mercaptoethanol. Eluted proteins were analyzed by western blotting.

### Statistics and reproducibility

GraphPad Prism (version 10.3.0) or R (version 4.5.1) were used to generate graphs and perform statistical analyses. Data were assessed for normal distribution, and appropriate statistical tests were applied accordingly. Statistical significance was determined using unpaired t-test for comparison of two samples and one-way ANOVA with multiple-comparison corrections for greater than two samples. All quantitative experiments were independently repeated at least three times with similar results.

## Data availability

Atomic coordinates and structure factors have been deposited in the Protein Data Bank under accession numbers XXXX (SPD^VDVD^) and XXXX (mini-LT-C:SPD^VDVD^).

## Supplementary information

This article contains supporting information.

## Acknowledgements

We thank Dr. Barry Milavetz (University of North Dakota) for the gift of the SV40 genome, and Dr. Bernard Moss for his gift of the BS-C-1 cells.

## Author contributions

Conceptualization, M.J.S. and Y.J.M.; Investigation, A.L.W., S.D., Y.M., J.R.A., S.P., M.E.C., A.T.Q.C., C.P.J., K.I., L.M.J., M.J.S., and Y.J.M.; Writing – Original Draft, A.L.W., S.D., Y.M., L.M.J., M.J.S., and Y.J.M.; Writing – Review & Editing, A.L.W., S.D., Y.M., L.M.J., M.J.S., and Y.J.M.; Funding Acquisition, Y.J.M.; Supervision, M.J.S. and Y.J.M.

## Funding and additional information

This study was supported by the National Institutes of Health (R01 CA233700 to Y.J.M. and M.J.S.) and the Intramural Research Program of the NIH, National Cancer Institute, Center for Cancer Research (ZIA BC 012086 to Y.J.M.). This work is based upon research conducted at the Northeastern Collaborative Access Team beamlines, which are funded by the National Institute of General Medical Sciences from the National Institutes of Health (P30 GM124165) and NIH-ORIP HEI grant (S10OD021527). This research used resources of the Advanced Photon Source, a U.S. Department of Energy (DOE) Office of Science User Facility operated for the DOE Office of Science by Argonne National Laboratory under Contract No. DE-AC02-06CH11357.

The contributions of the NIH author(s) were made as part of their official duties as NIH federal employees, are in compliance with agency policy requirements, and are considered Works of the United States Government. However, the findings and conclusions presented in this paper are those of the author(s) and do not necessarily reflect the views of the NIH or the U.S. Department of Health and Human Services.

---

**Supplementary Figure 1.**
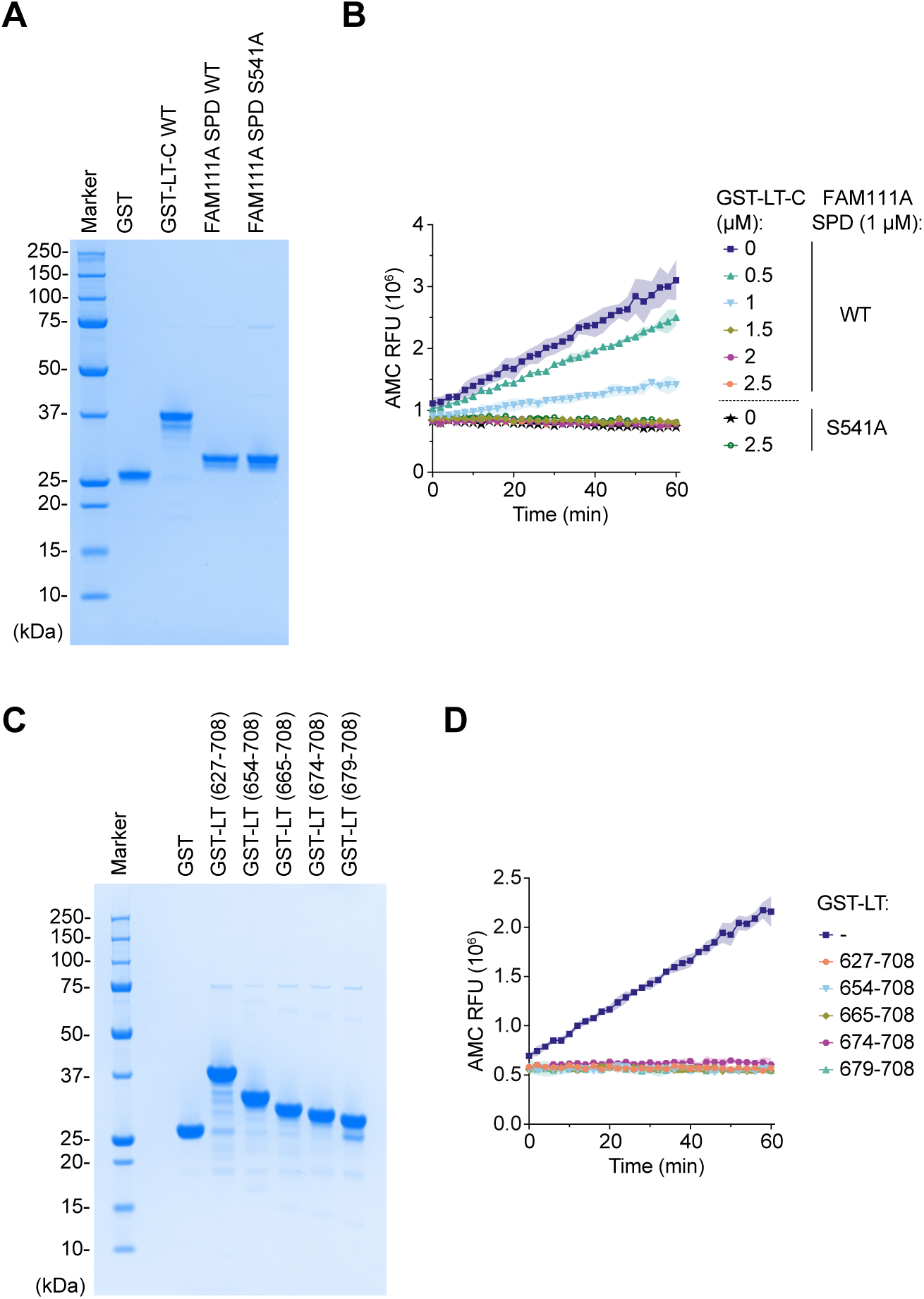
Purification and activity of recombinant FAM111A SPD and GST-LT-C proteins. **(A)** One microgram of purified recombinant GST, GST-LT-C (WT), FAM111A serine protease domain (SPD; WT) and FAM111A SPD (S541A) analyzed by SDS-PAGE and visualized by Coomassie staining. Molecular weight markers are shown on the left. **(B)** Time course of *in vitro* FAM111A SPD activity in the presence of increasing concentrations of GST-LT-C. Reactions without GST-LT-C contain GST as a control. Peptidase activity was measured using Suc-AAPF-AMC as a substrate and fluorescence from released AMC was measured over time for 1 hr at 37°C. Values represent the mean of three technical replicates, and standard deviation is indicated by the shaded area. RFU: relative fluorescence units. **(C)** Two micrograms of purified recombinant GST and the indicated GST-LT proteins analyzed by SDS-PAGE. Molecular weight markers are shown on the left. **(D)** Time course of *in vitro* FAM111A SPD activity in the presence of GST-LT truncation proteins. Reactions without GST-LT contain GST as a control. Peptidase activity was measured as in (B).

**Supplementary Figure 2.**
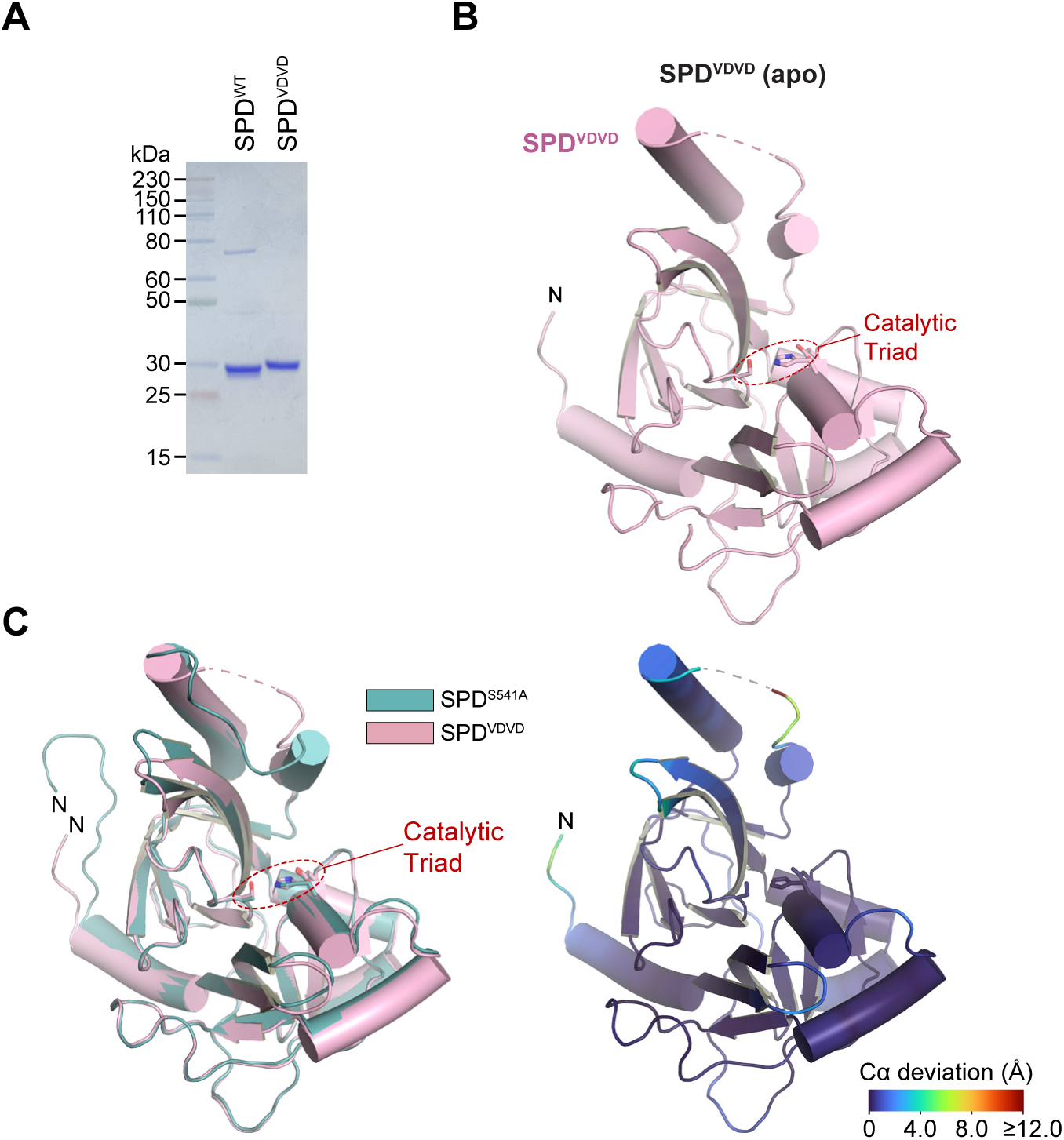
Purification and structural characterization of FAM111A SPD^VDVD^ (apo). **(A)** 2.4 micrograms of purified recombinant FAM111A serine protease domain (SPD^VDVD^; V347D/V351D) analyzed by SDS-PAGE and visualized by Coomassie staining. Molecular weight markers are shown on the left. **(B)** Overall structure of the apo SPD^VDVD^ crystal structure shown as a cartoon representation. The catalytic triad is highlighted. **(C)** Superposition of the apo SPD^VDVD^ (pink) and previously reported SPD^S541A^ (teal) structures. Left, structural overlay. Right, Cα deviation mapped onto the SPD^VDVD^ structure, with colors indicating the magnitude of local structural differences.

**Supplementary Figure 3.**
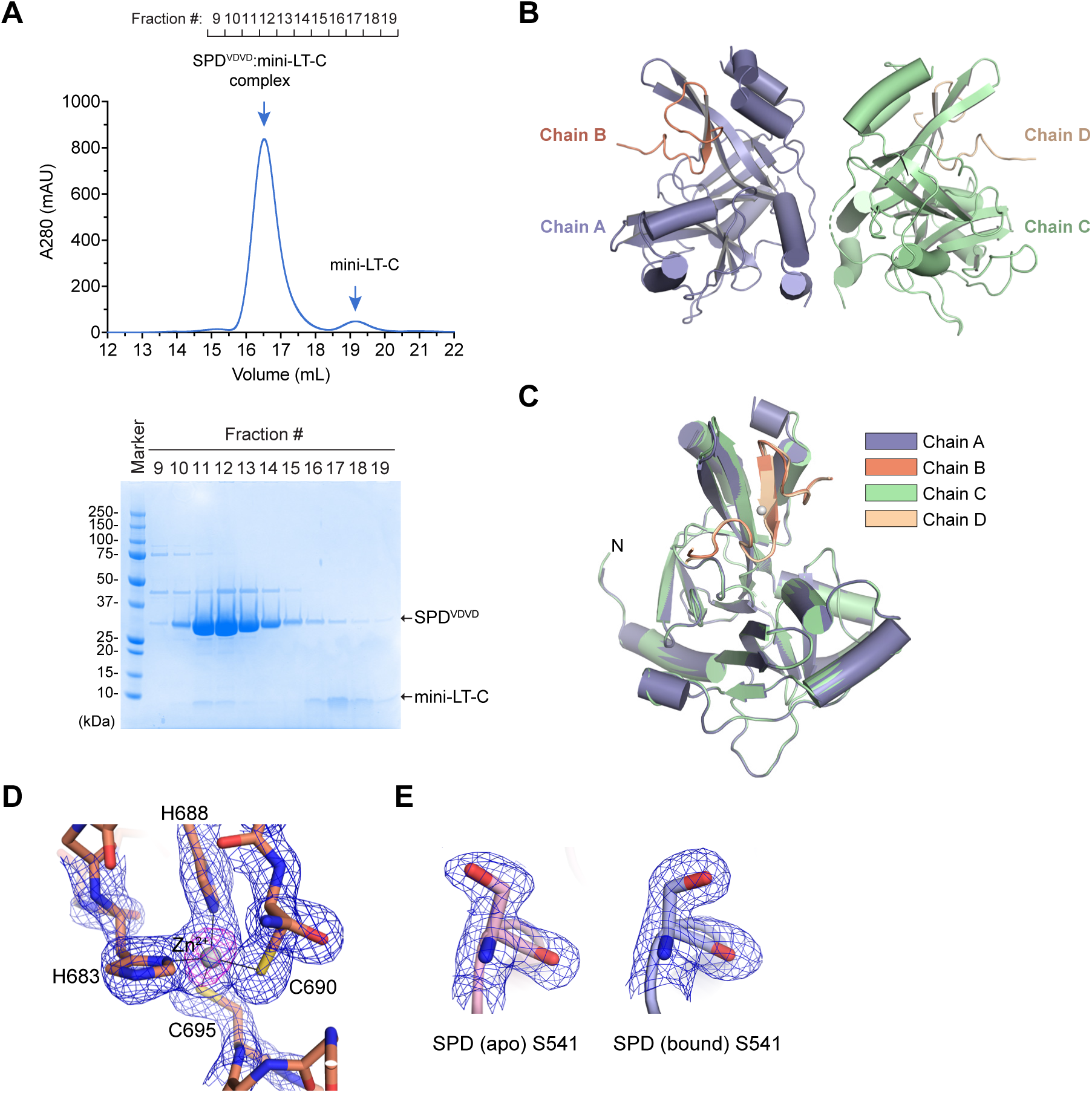
Purification and structural characterization of the SPD^VDVD^:mini-LT-C complex. **(A)** Purification of the SPD^VDVD^:mini-LT-C complex by size-exclusion chromatography. Top, elution profile showing the major protein peak used for crystallization. Bottom, Coomassie-stained SDS-PAGE analysis of the indicated fractions. The positions of SPD^VDVD^ and mini-LT-C are indicated on the right, and molecular weight markers are shown on the left. **(B)** Overall structure of the asymmetric unit of the SPD^VDVD^:mini-LT-C crystal. The asymmetric unit contains two SPD^VDVD^:mini-LT-C complexes (chains A/B and C/D). **(C)** Superposition of the two SPD^VDVD^:mini-LT-C complexes in the asymmetric unit. Chains are colored as indicated. **(D)** 2mFo-DFc electron density map (blue mesh, contoured at 1σ) and anomalous difference map (magenta mesh, contoured at 5σ) depicting the zinc-binding site of mini-LT-C. The zinc ion and coordinating residues His683, His688, Cys690, and Cys695 are shown as sticks. **(E)** 2mFo-DFc electron density map contoured at 1σ showing density for the catalytic Ser541 in the apo SPD^VDVD^ structure (left) and the SPD^VDVD^:mini-LT-C complex (right).

**Supplementary Figure 4.**
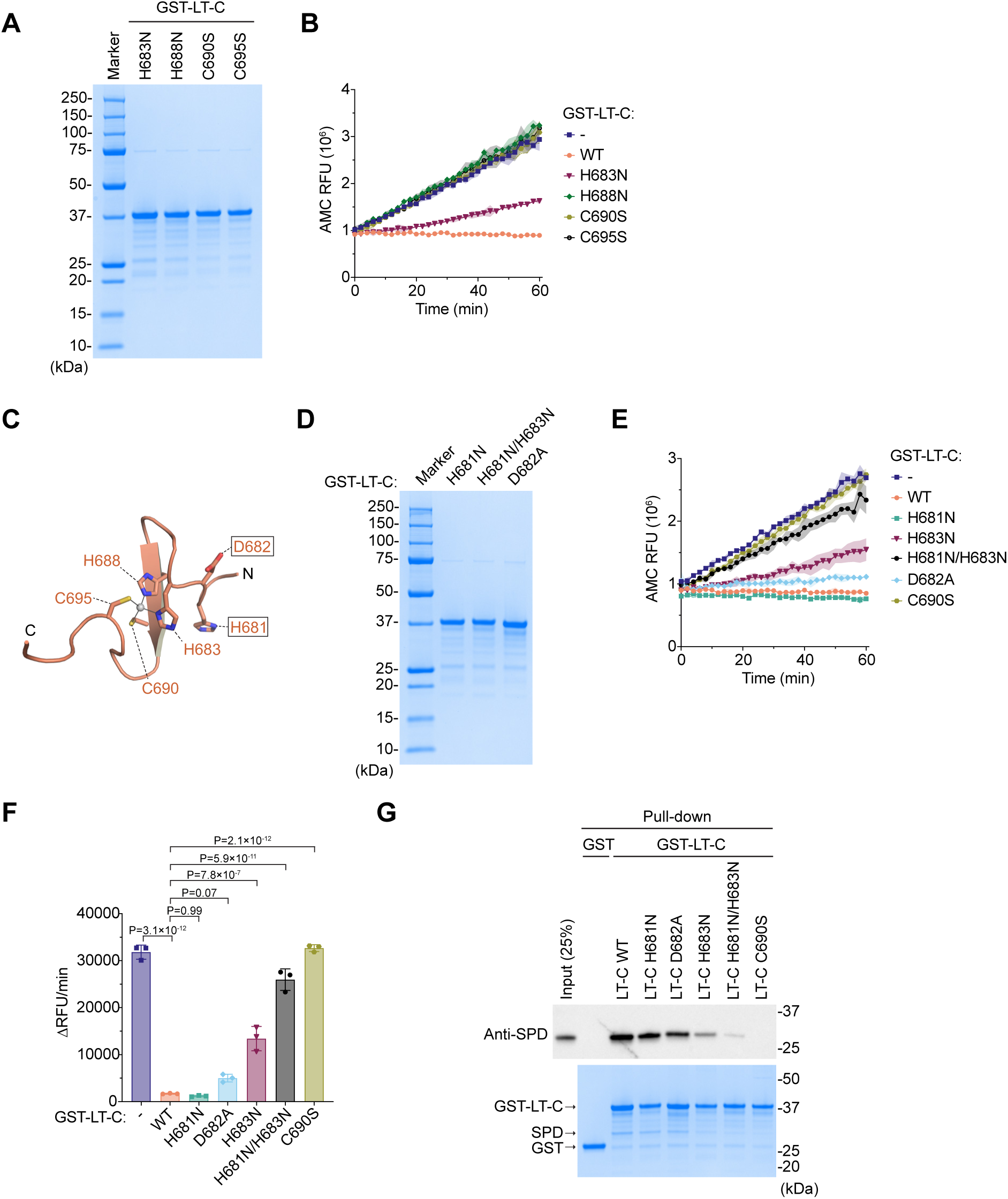
Effects of zinc-binding motif mutations on LT-C-mediated inhibition of and binding to FAM111A. **(A)** One microgram of purified recombinant GST-LT-C zinc-binding motif mutants (H683N, H688N, C690S, and C695S) analyzed by SDS-PAGE and visualized by Coomassie staining. Molecular weight markers are shown on the left. **(B)** Time course of *in vitro* FAM111A SPD activity in the presence of GST-LT-C (wild-type or zinc-binding motif mutants H683N, H688N, C690S, and C695S). Reactions without GST-LT-C contain GST as a control. Peptidase activity was measured using Suc-AAPF-AMC as a substrate and fluorescence from released AMC was measured over time for 1 hr at 37°C. Values represent the mean of three technical replicates, and standard deviation is indicated by the shaded area. RFU: relative fluorescence units. **(C)** Close-up view of the LT-C zinc-binding motif. Side chains of residues H681 and D682, in addition to the zinc-coordinating residues (H683, H688, C690, and C695) are shown as sticks. **(D)** One microgram of purified recombinant GST-LT-C mutants (H681N, H681N/H683N, and D682A) analyzed by SDS-PAGE and visualized by Coomassie staining. **(E)** Time course of in vitro FAM111A SPD activity in the presence of GST-LT-C, either wild-type or mutants (H681N, H681N/H683N, and D682A). Reactions without GST-LT-C contain GST as a control. Peptidase activity was measured as in (B). Values represent the mean of three technical replicates, and standard deviation is indicated by the shaded area. **(F)** In vitro protease activity of FAM111A SPD measured in the presence of GST-LT-C (wild-type or mutants H681N, H683N, H681N/H683N, D682A, and C690S). Reactions without GST-LT-C contain GST as a control. Values are mean ± SD of three technical replicates. Statistical significance was determined by one-way ANOVA with Dunnett’s multiple-comparison test. **(G)** In vitro GST pull-down assay of FAM111A SPD with GST-LT-C (wild-type, or mutants H681N, H683N, H681N/H683N, D682A, and C690S). Bound FAM111A SPD was detected by western blotting (top), and GST proteins were visualized by Coomassie staining (bottom). Molecular weight markers are shown on the right.

**Supplementary Figure 5.**
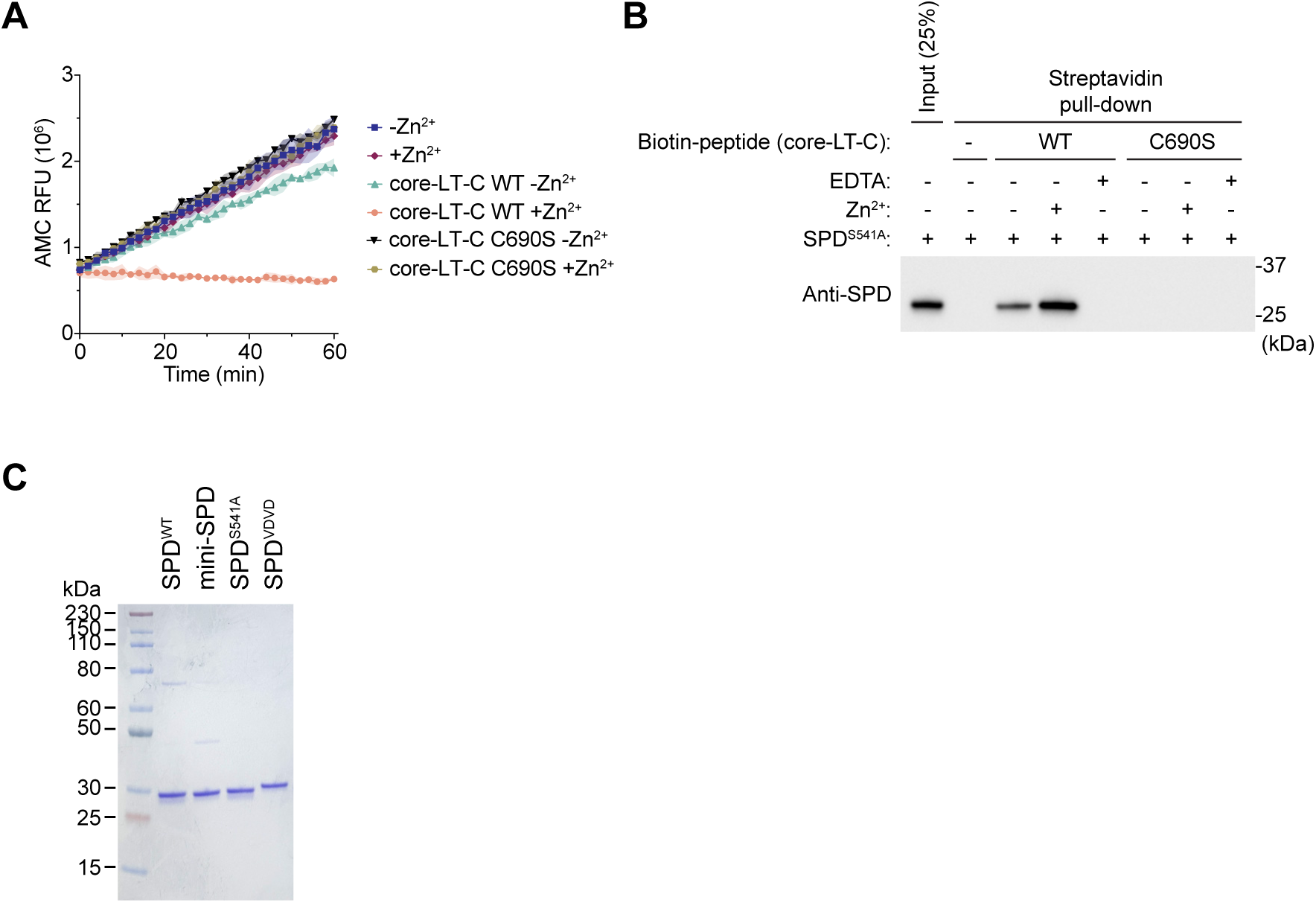
Zinc is required for LT-C-mediated inhibition and interaction with FAM111A. **(A)** Time course of *in vitro* FAM111A SPD activity measured in the presence of a synthetic LT-C peptide corresponding to residues 679-704 (wild-type or C690S) incubated with or without ZnCl_2_. Peptidase activity was measured using Suc-AAPF-AMC as a substrate and fluorescence from released AMC was measured over time for 1 hr at 37°C. Values represent the mean of three technical replicates, and standard deviation is indicated by the shaded area. RFU: relative fluorescence units. **(B)** *In vitro* streptavidin pull-down assay using biotinylated LT-C peptides corresponding to residues 679-704 (wild-type or C690S) incubated with FAM111A SPD S541A in the presence or absence of a twofold molar excess of ZnCl_2_. Where indicated, EDTA was added to the reaction. Bound FAM111A SPD was detected by western blotting. Molecular weight markers are shown on the right. **(C)** The indicated purified recombinant SPD proteins (1.2 µg each) were analyzed by SDS-PAGE and visualized by Coomassie staining. Molecular weight markers are shown on the left.

**Supplementary Figure 6.**
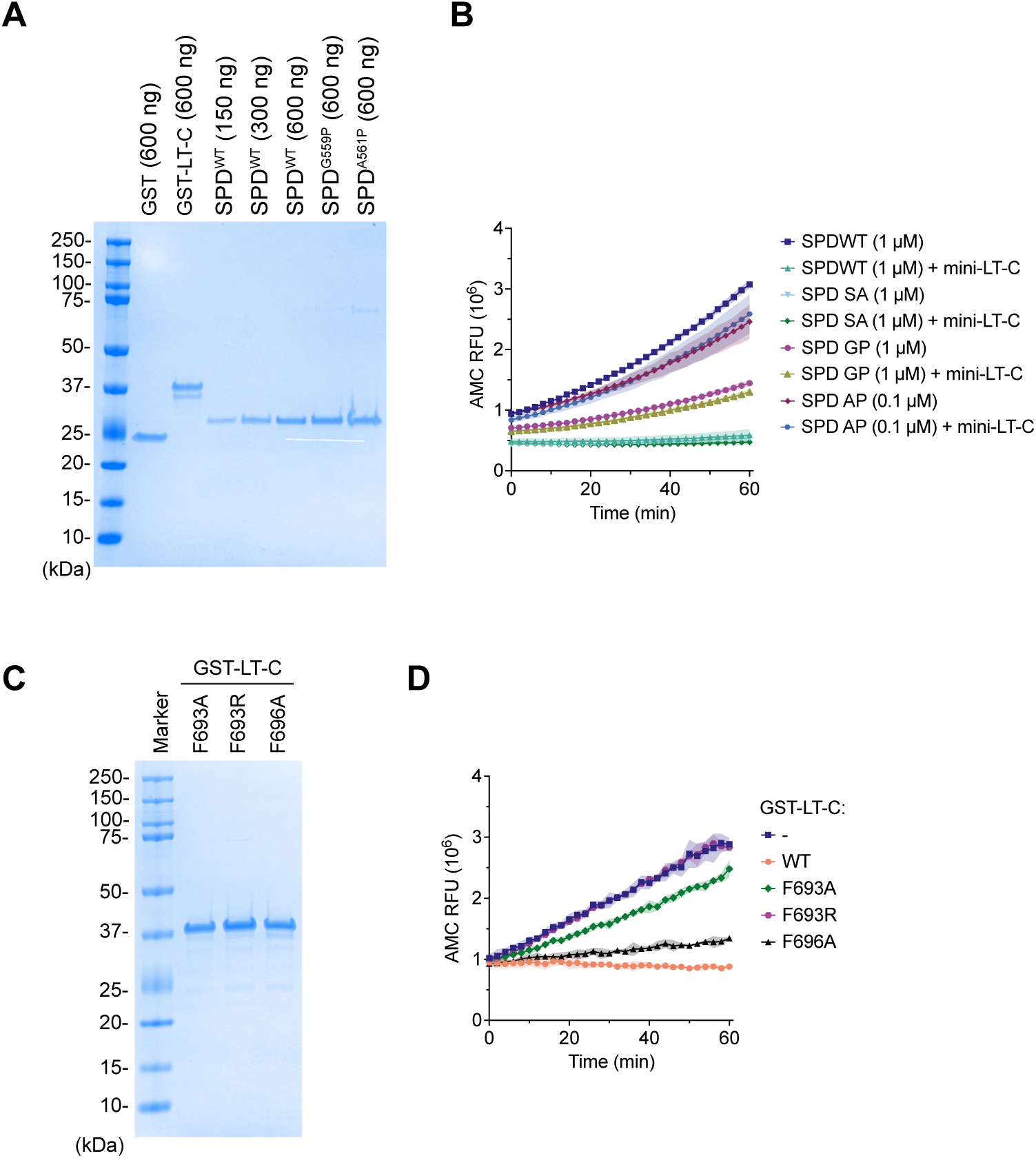
Contributions of the β10 interface and the P1-like phenylalanine residue to LT-C-mediated FAM111A inhibition. **(A)** The indicated amounts of purified recombinant GST, GST-LT-C, and SPD (wild-type, G559P, and A561P) proteins analyzed by SDS-PAGE and visualized by Coomassie staining. Molecular weight markers are shown on the left. **(B)** Time course of *in vitro* activity of FAM111A SPD (wild-type, G559P, and A561P) in the absence or presence of mini-LT-C. Peptidase activity was measured using Suc-AAPF-AMC as a substrate and fluorescence from released AMC was measured over time for 1 hr at 37°C. Values represent the mean of three technical replicates, and standard deviation is indicated by the shaded area. RFU: relative fluorescence units. **(C)** One microgram of purified recombinant GST-LT-C mutants (F693A, F693R, and F696A) analyzed by SDS-PAGE and visualized by Coomassie staining. Molecular weight markers are shown on the left. **(D)** Time course of *in vitro* FAM111A SPD activity in the presence of GST-LT-C (wild-type or phenylalanine mutants F693A, F693R, and F696A). Reactions without GST-LT-C contain GST as a control. Peptidase activity was measured as in (B). Values represent the mean of three technical replicates, and standard deviation is indicated by the shaded area.

**Supplementary Figure 7.**
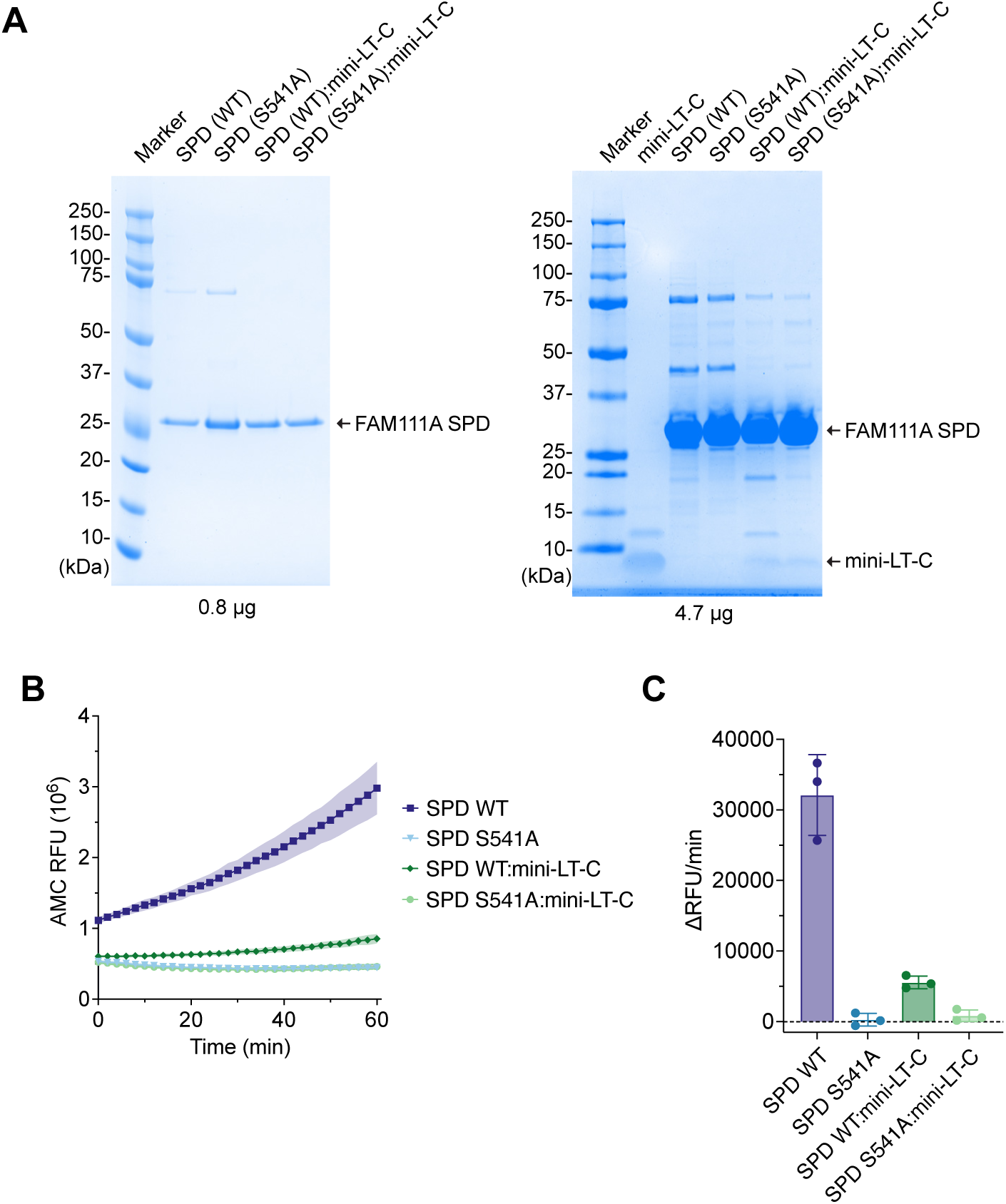
Purification and peptidase activity of SPD:mini-LT-C complexes. **(A)** SDS-PAGE analysis of purified mini-LT-C, FAM111A SPD (wild-type and S541A), and size-exclusion chromatography-purified complexes of SPD (wild-type or S541A) with mini-LT-C. Molecular weight markers are shown on the left. **(B)** Time course of FAM111A SPD activity for SPD alone (wild-type) and for purified SPD-mini-LT-C complexes (wild-type or S541A). Peptidase activity was measured using Suc-AAPF-AMC as a substrate and fluorescence from released AMC was measured over time for 1 hr at 37°C. Values represent the mean of three technical replicates, and standard deviation is indicated by the shaded area. RFU: relative fluorescence units. **(C)** Peptidase activity of SPD expressed as change in relative fluorescence units per minute (ΔRFU/min). Values are mean ± SD of three technical replicates.

**Supplementary Figure 8.**
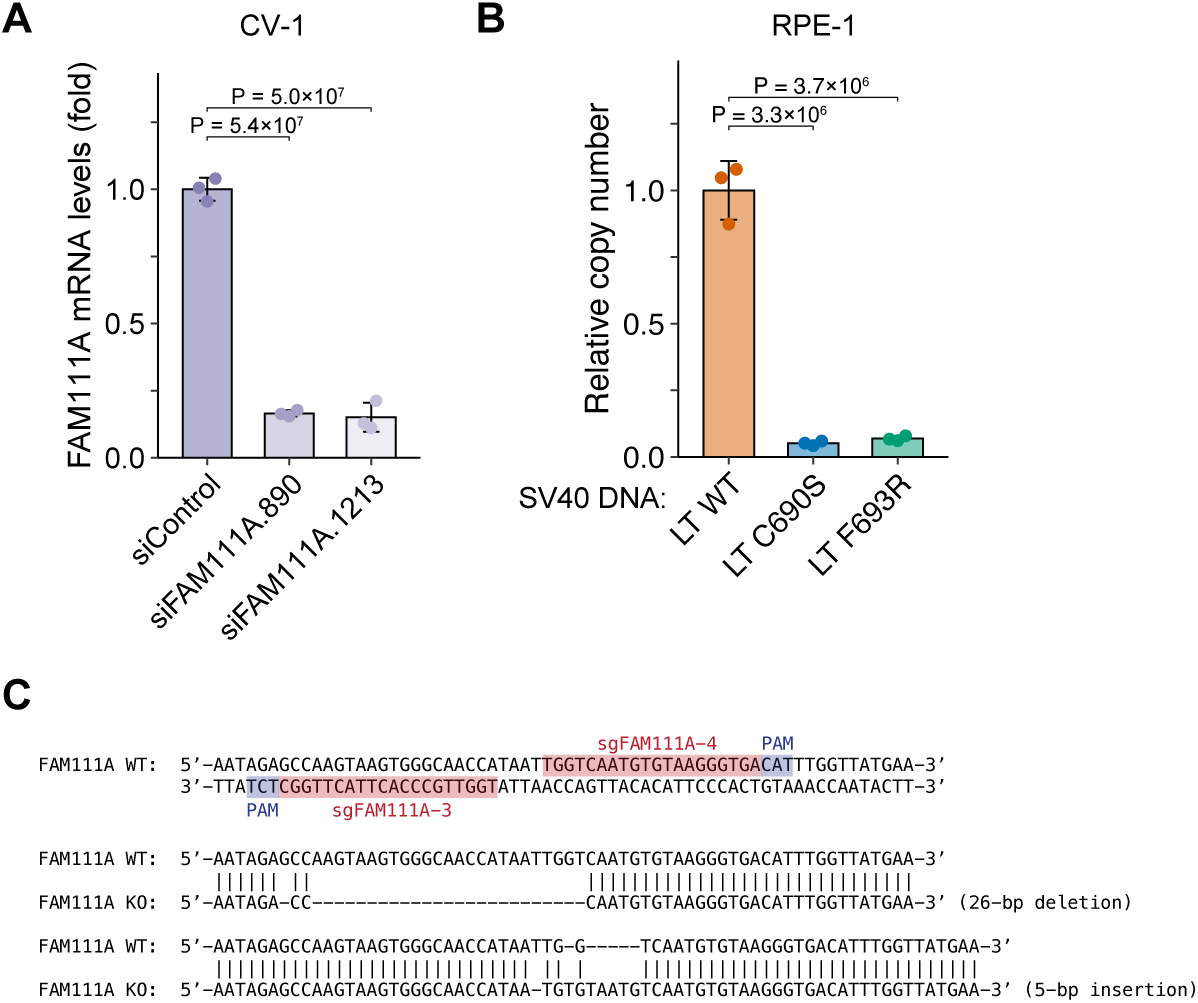
Validation of FAM111A knockdown and knockout. **(A)** Relative FAM111A mRNA levels in CV-1 cells transfected with control siRNA (siControl) or two independent siRNAs targeting FAM111A. mRNA levels were measured by quantitative PCR, normalized to GAPDH, and expressed relative to the control sample. Data represent mean ± SD of three biological replicates. Statistical significance was determined by one-way ANOVA with Dunnett’s multiple-comparison test. **(B)** Quantification of SV40 genome copy number in RPE-1 cells transfected with WT, C690S, F693R SV40 genomic DNA. Viral genomic copy number was determined by quantitative PCR 72 hr after transfection. Values are mean ± SD of three technical replicates. Statistical significance was determined by one-way ANOVA with Dunnett’s multiple-comparison test. **(C)** Schematic of sgRNA target sites within FAM111A exon 4 showing the positions of sgFAM111A-3 and sgFAM111A-4 and their PAM sequences. Sequence alignments of wild-type (WT) and knockout (KO) alleles are shown. Sizes of each insertion and deletion at the target sites are indicated.

**Supplementary Table 1.**
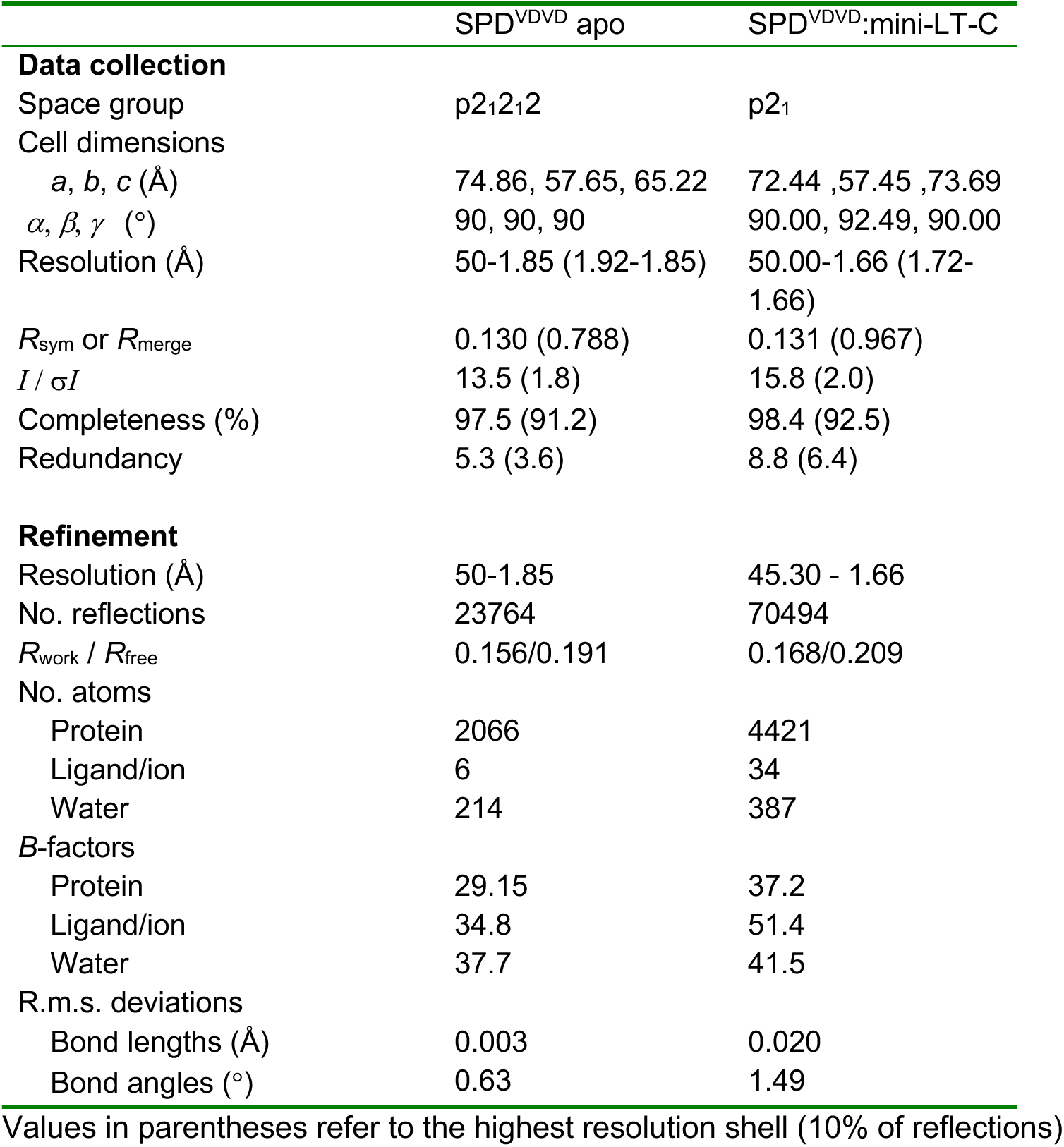
X-ray crystallographic data collection and refinement statistics.

**Supplementary Table 2.**
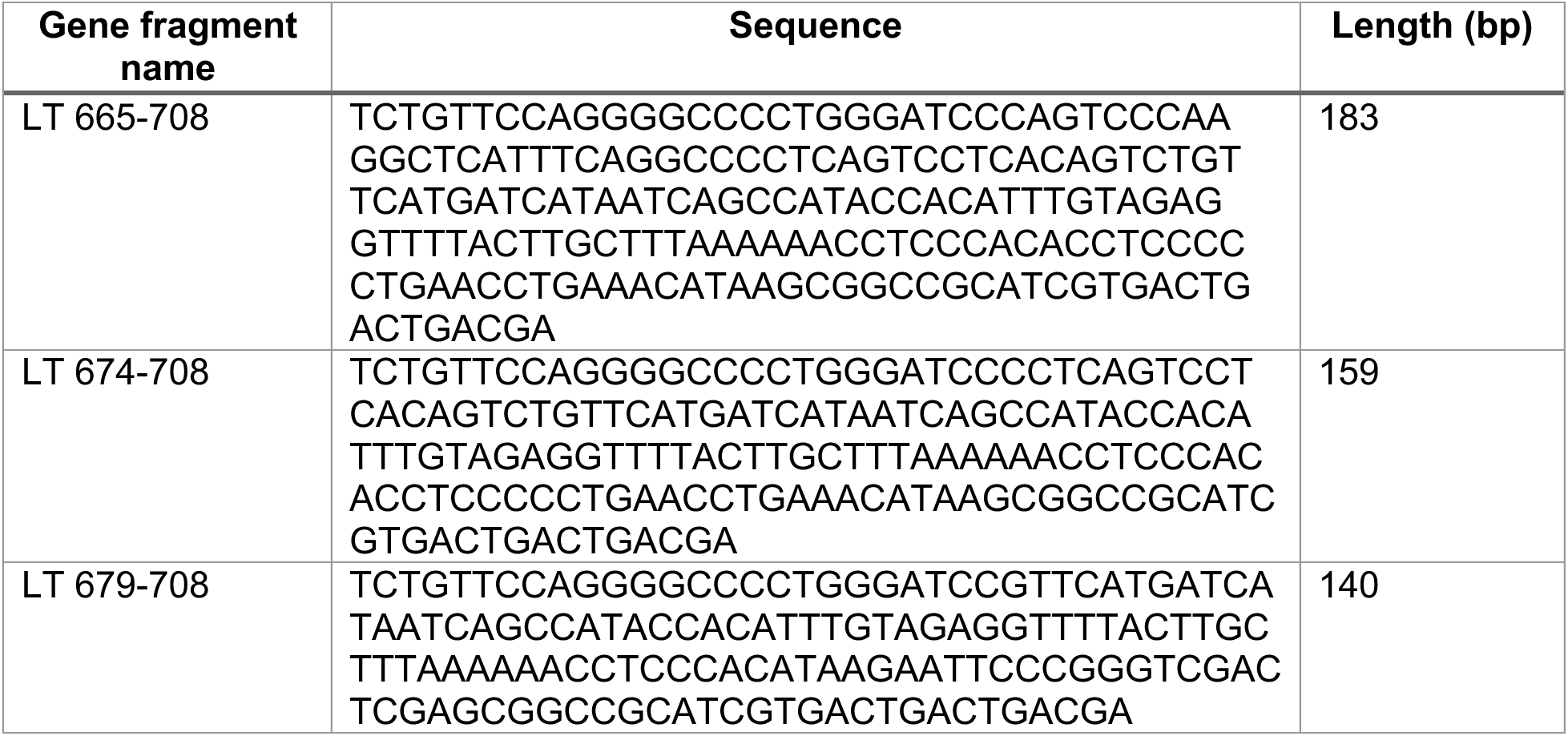
Gene fragments used in this study.

**Supplementary Table 3.**
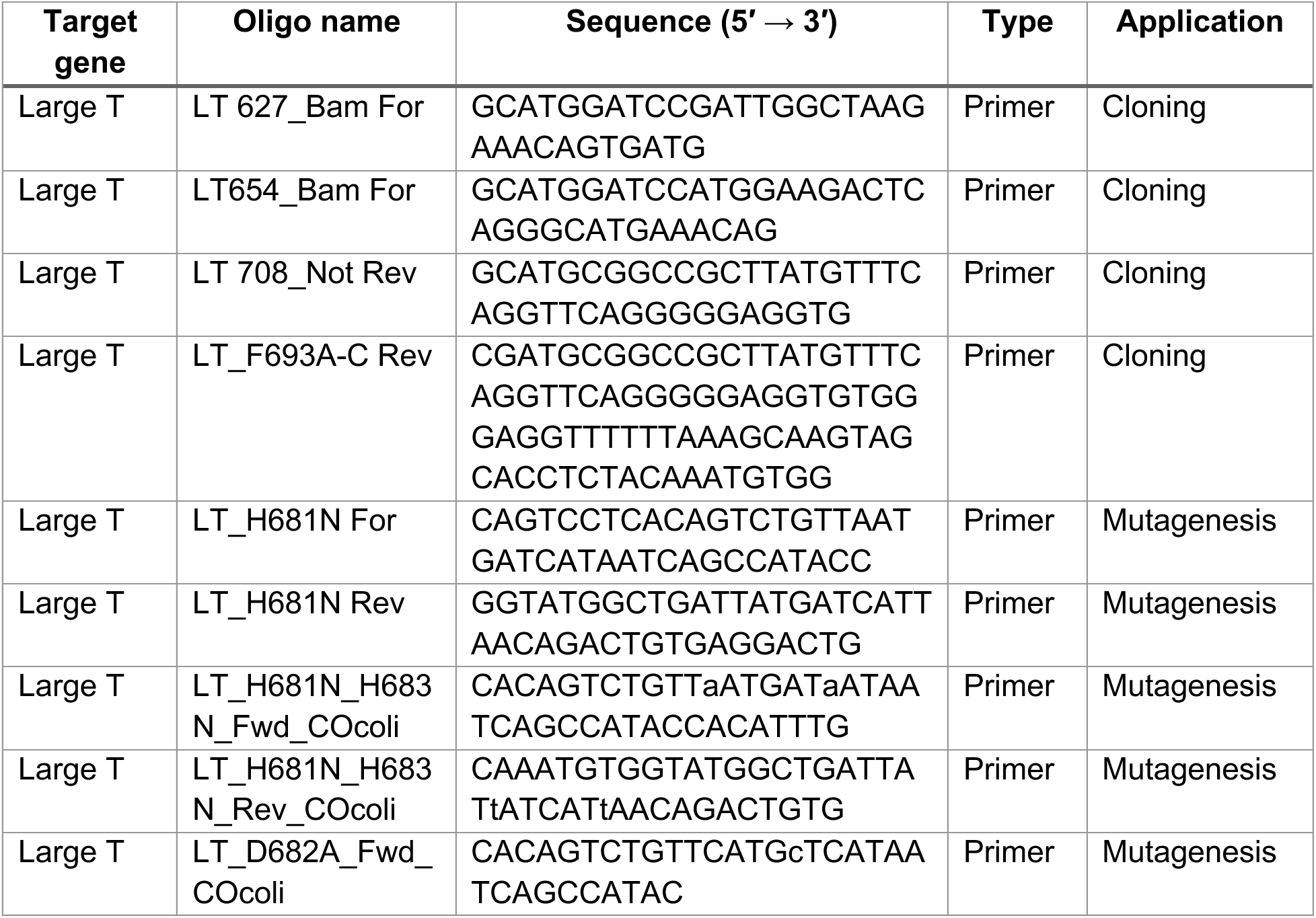
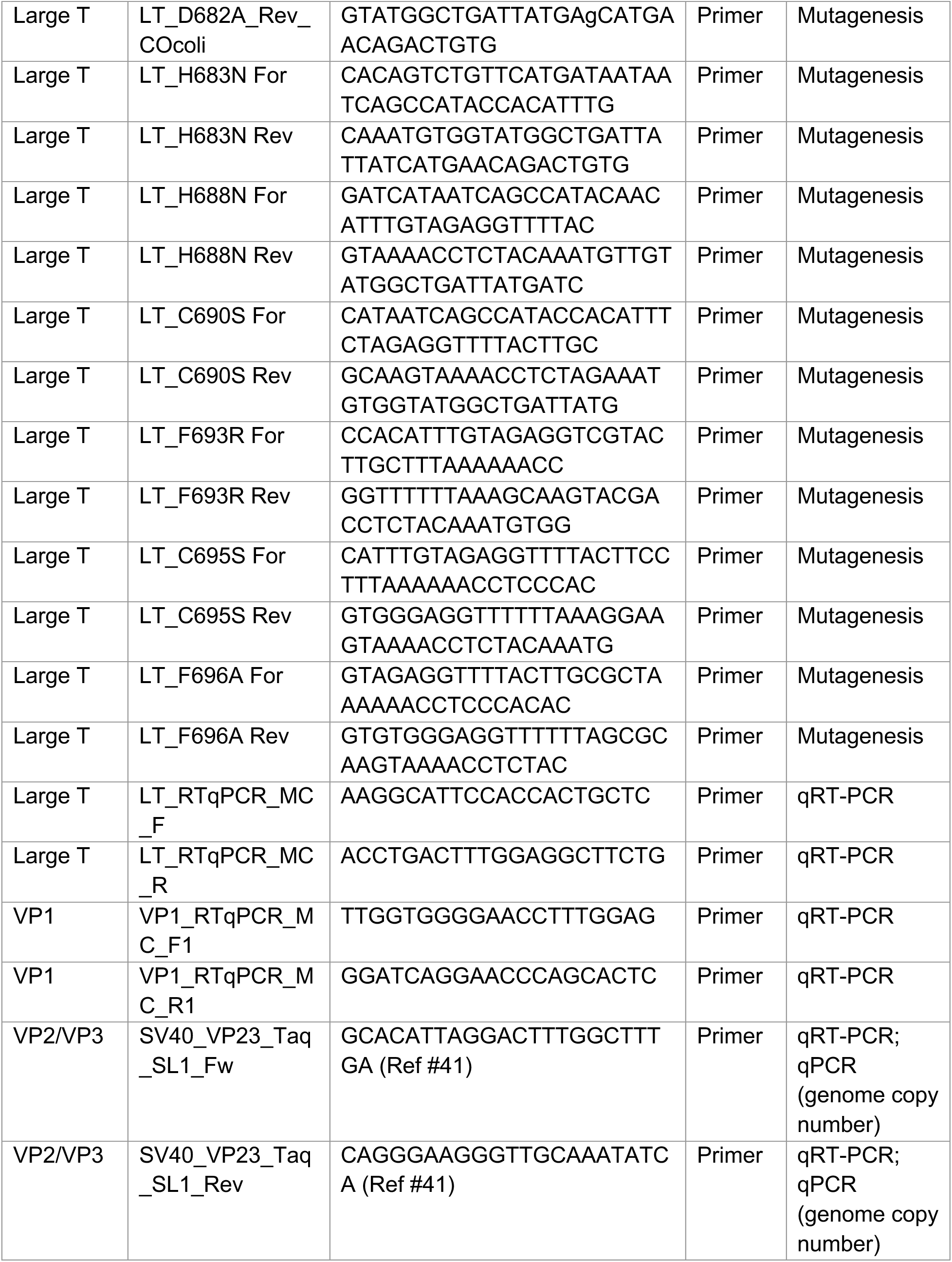
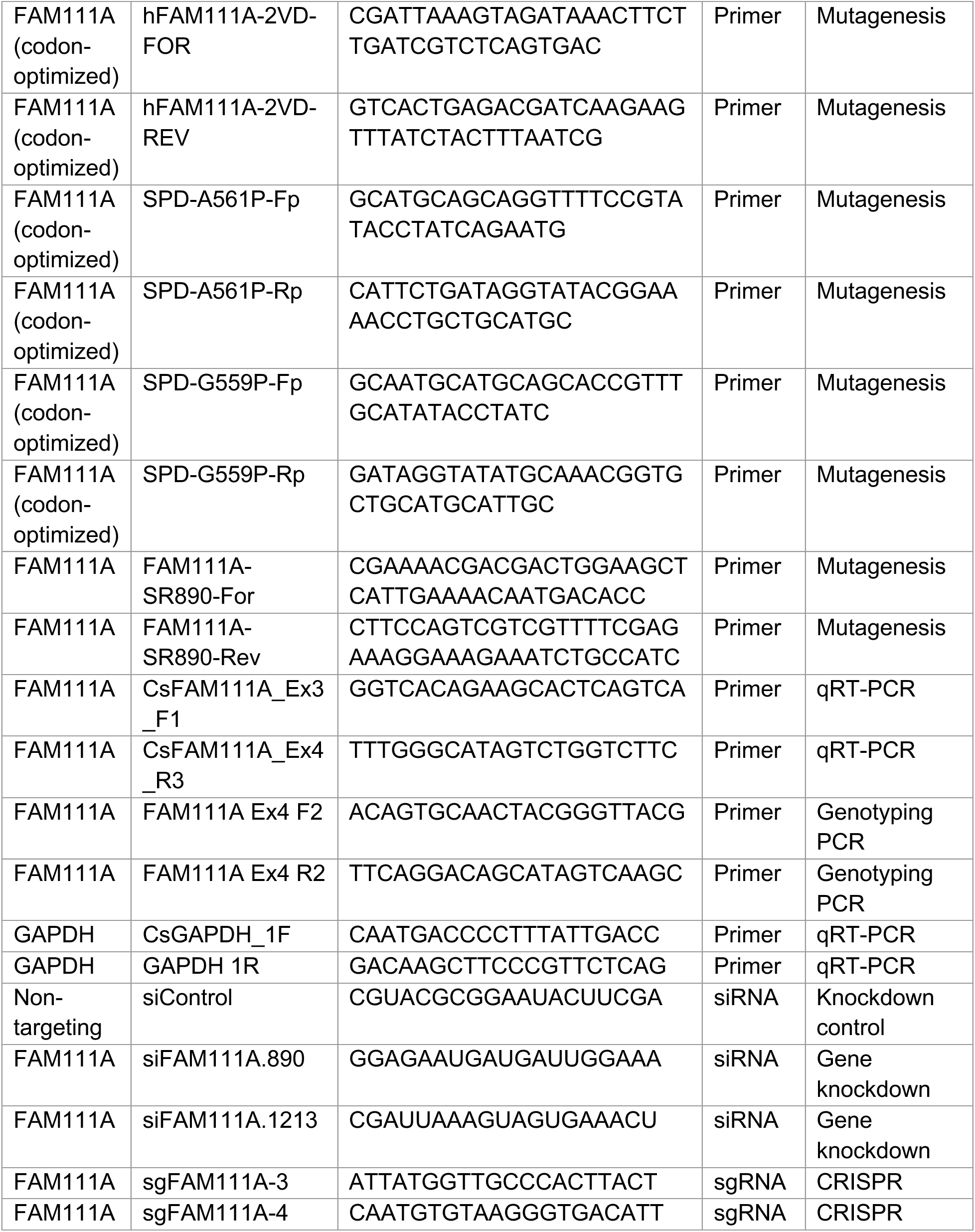
Oligonucleotides and guide RNAs used in this study.

**Supplementary Table 4.** Plasmids used in this study.

| **Plasmid name** | **Backbone** | **Insert / description** | **Tag** | **Source** |
| --- | --- | --- | --- | --- |
| pDB.His.MBP/FAM111A SPD (WT; codon-optimized) | pDB.His.MBP | Human FAM111A SPD (aa 335-611) | His<sub>6</sub>-MBP | Ref #22 |
| pDB.His.MBP/FAM111A SPD (S541A; codon-optimized) | pDB.His.MBP | Catalytically inactive FAM111A SPD (S541A) | His<sub>6</sub>-MBP | Ref #22 |
| pDB.His.MBP/FAM111A mini-SPD (codon-optimized) | pDB.His.MBP | Dimerization site mutant of FAM111A SPD | His<sub>6</sub>-MBP | Ref #22 |
| pDB.His.MBP/FAM111A SPD (VDVD; codon-optimized) | pDB.His.MBP | Dimerization site mutant of FAM111A SPD | His<sub>6</sub>-MBP | Ref #22 |
| pDB.His.MBP/FAM111A SPD (G556P; codon-optimized) | pDB.His.MBP | β10 mutant of FAM111A SPD | His<sub>6</sub>-MBP | This study |
| pDB.His.MBP/FAM111A SPD (A561P; codon-optimized) | pDB.His.MBP | β10 mutant of FAM111A SPD | His<sub>6</sub>-MBP | This study |
| pGEX-6P-2/LT-C (WT) | pGEX-6P-2 | SV40 Large T C-terminal region (aa 627-708) | GST | This study |
| pGEX-6P-2/LT-C (H681N) | pGEX-6P-2 | LT-C H681N mutant | GST | This study |
| pGEX-6P-2/LT-C (H681N/H683N) | pGEX-6P-2 | LT-C double mutant H681N/H683N | GST | This study |
| pGEX-6P-2/LT-C (D682A) | pGEX-6P-2 | LT-C D682A mutant | GST | This study |
| pGEX-6P-2/LT-C (H683N) | pGEX-6P-2 | LT-C H683N mutant | GST | This study |
| pGEX-6P-2/LT-C (H688N) | pGEX-6P-2 | LT-C H688N mutant | GST | This study |
| pGEX-6P-2/LT-C (C690S) | pGEX-6P-2 | LT-C C690S mutant | GST | This study |
| pGEX-6P-2/LT-C (F693A) | pGEX-6P-2 | LT-C F693A mutant | GST | This study |
| pGEX-6P-2/LT-C (F693R) | pGEX-6P-2 | LT-C F693R mutant | GST | This study |
| pGEX-6P-2/LT-C (C695S) | pGEX-6P-2 | LT-C C695S mutant | GST | This study |
| pGEX-6P-2/LT-C (F696A) | pGEX-6P-2 | LT-C F696A mutant | GST | This study |
| pGEX-6P-2/LT (654-708) | pGEX-6P-2 | SV40 Large T (aa 654-708) | GST | This study |
| pGEX-6P-2/LT (665-708) | pGEX-6P-2 | SV40 Large T (aa 665-708) | GST | This study |
| pGEX-6P-2/LT (674-708) | pGEX-6P-2 | SV40 Large T (aa 674-708) | GST | This study |
| pGEX-6P-2/mini-LT-C (WT) | pGEX-6P-2 | SV40 Large T (aa 679-708) | GST | This study |
| pUC18/SV40 (WT) | pUC18 | SV40 genome (strain 776) | N/A | This study |
| pUC18/SV40 (C690S) | pUC18 | SV40 genome encoding LT C690S | N/A | This study |
| pUC18/SV40 (F693R) | pUC18 | SV40 genome encoding LT F693R | N/A | This study |
| pLVX2-IRES-Puro/<br>FAM111A SR890 (WT) | pLVX2-IRES-Puro | Full-length human FAM111A (resistant to siFAM111A.890) | N/A | This study |
| pLVX2-IRES-Puro/<br>FAM111A SR890 (S541A) | pLVX2-IRES-Puro | Catalytically inactive FAM111A (resistant to siFAM111A.890) | N/A | This study |
| pLVX2-IRES-Puro/<br>FAM111A SR890 (YFAA) | pLVX2-IRES-Puro | PIP motif mutant (YFAA) (resistant to siFAM111A.890) | N/A | This study |
| pMD2.G | N/A | Lentiviral packaging plasmid | N/A | Addgene #12259 |
| psPAX2 | N/A | Lentiviral envelope plasmid | N/A | Addgene #12260 |

**Supplementary Table 5.**
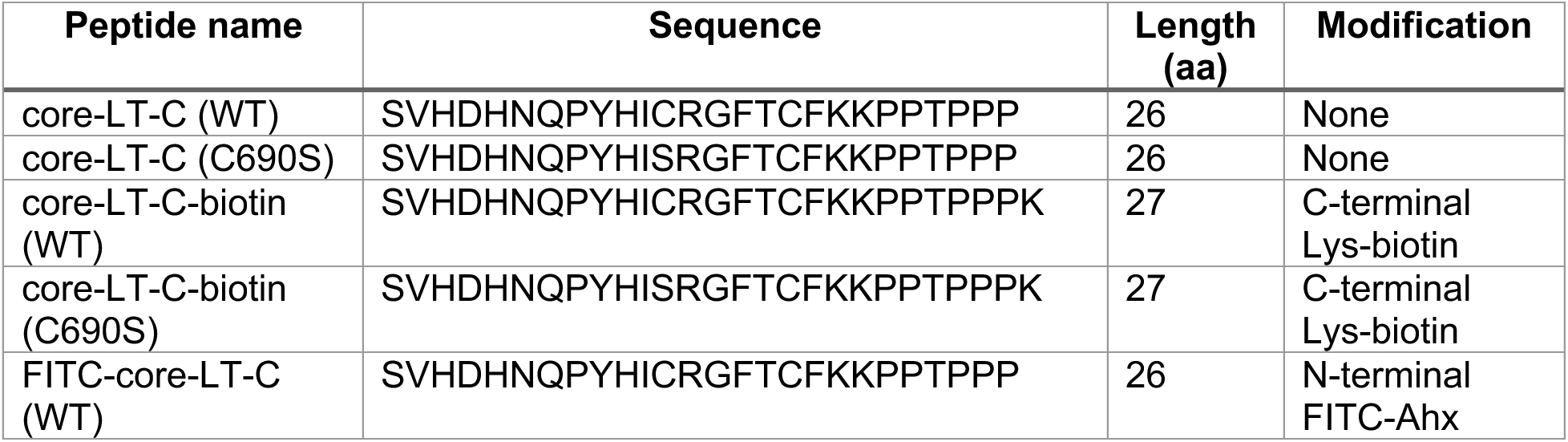
Synthetic peptides used in this study.

**Supplementary Table 6.** Antibodies used in this study.

| Target protein | Application | Host species | Source | Catalog number | Dilution for western blot |
| --- | --- | --- | --- | --- | --- |
| FAM111A | Western blot | Rabbit | Abcam | ab184572 | 1:1,000 |
| FAM111A | Western blot | Rabbit | Sigma | HPA040176 | 1:1,000 |
| FAM111A (SPD) | Western blot | Rabbit | Cocalico (custom) | N/A | 1:1,000 |
| SV40 Large T antigen | Western blot, Immunoprecipitation | Mouse | Abcam | ab16879 | 1:2,000 |
| SV40 VP1 | Western blot | Rabbit | Abcam | ab53977 | 1:2,000 |
| SV40 VP2/3 | Western blot | Rabbit | Abcam | ab53983 | 1:1,000 |
| GAPDH | Western blot | Mouse | GeneTex | GTX627408 | 1:1,000 |

